# Spatially coordinated cell cycle activity and motility govern mammary ductal growth and tip bifurcation

**DOI:** 10.1101/2022.08.29.505725

**Authors:** SM Myllymäki, B Kaczynska, Q Lan, ML Mikkola

## Abstract

Branching morphogenesis is the common evolutionary solution of multiple organs to combine maximal epithelial function with compact organ size. It involves successive rounds of branch elongation and branch point generation to generate a branched epithelial network. Branch points form most commonly at the tips of branches as they split into two. However, it is unclear how epithelial cells in tips drive both elongation and branching. In this study, we used ex vivo live imaging to identify these fundamental cellular mechanisms in the embryonic mammary gland. Our 4D analyses show that tips of branches are driven forward by directional cell migration, while elongation of the subtending duct is supported by cell proliferation that feeds a retrograde flow of lagging cells into the duct, established upon differential cell motility. Tip bifurcation involved localized repression of both cell cycle and cell motility at the branch point. Cells in the nascent daughter tips remained proliferative, but changed the direction of their movement to elongate new branches. Our study also reports the fundamental importance of the contractile actin cytoskeleton in regulating branch point generation in the mammary epithelium.

## Introduction

Branching morphogenesis is a conserved developmental program that molds an epithelial tube or chord into the form of a branched tree. As a result, the functional surface area of the epithelium is increased manifold – a principle adopted by the lungs, kidneys, and many secretory organs, including the mammary gland. During development, each organ follows a unique pattern of epithelial growth and branching, orchestrated by inductive signals from the surrounding mesenchymal tissue (Lang et al. 2021). Inductive signals elicit changes in epithelial cell behaviors leading to elongation of branches and generation of new ones, fundamental to the construction of branched organ structure. In the mammary gland, these crucial cellular mechanisms still await recognition. Beyond secreting signaling molecules, the contribution of mesenchymal cells to these activities, is also not fully understood. Epithelial cells are self-sufficient in forming branching structures in mesenchyme-free 3D organoid culture when supplemented with appropriate growth factors, although the pattern of branching is not organotypic (Nerger & Nelson 2019).

Two topological domains of the growing epithelium can serve as branch points, from which new branches arise in vivo. Branches can sprout from the side of an existing branch, which is referred to as lateral or side-branching, but this has only been documented in the lung and mammary gland (Iber & Menshykau 2013). In all branching organs, the blunt ended tips of branches can split into two or more branches – a process called terminal bifurcation or clefting. In the lung and kidney, the tip transforms from T- to Y-shape (Watanabe & Costantini 2004, Schnatwinkel & Niswander 2014), while in the salivary gland, it remains round with a cleft inserted (Harunaga et al. 2011). Between bifurcations, the tips invade into the mesenchymal tissue thereby elongating the stalk of the branch, also called “trailing duct”. The extent of elongation depends on the organ – in the salivary gland for example, the branch widens at the tip rather than elongates before splitting into new branches. In the mammary gland, branches elongate extensively between branch points during pubertal development (Scheele et al. 2017).

Mammary gland is a unique arborized organ as branching morphogenesis takes place in three developmental stages – embryonic, pubertal, and reproductive. In mice, branching morphogenesis begins when a sprout rooted to the embryonic surface ectoderm is induced to branch around E16 and yields a rudimentary tree of 10-20 branches by birth (E19-20) (Macias & Hinck 2012, Lindström et al. 2022). Lumen formation commences shortly before birth, and soon after, the final, bilayered structure of mammary ducts is established (Hogg et al., 1983). The inner layer of cells enclosing the lumen and outer layer of cells separated from the mesenchyme by a basement membrane, begin to segregate into distinct luminal and basal lineages already during late embryogenesis (Lilja et al. 2018). A second round of branching morphogenesis occurs during puberty when hormones instigate a massive expansion of the arborized ductal system over the fat pad, the stroma of the adult gland (Macias & Hinck 2012). At the onset of puberty, branch tips transform into bulbous terminal end buds (TEBs) containing multiple layers of luminal cells surrounded by a single layer of basal cells (Paine & Lewis 2017). TEBs are highly proliferative and thereby fuel ductal elongation by producing building blocks for the subtending duct. Morphometric analyses and computational modeling suggest that during puberty, new branches form mainly by TEB bifurcation (Scheele et al. 2017). In the adult gland, reproductive hormones stimulate side-branching during the estrous cycle and pregnancy (Brisken et al. 1998).

Limited mechanistic insight into branching morphogenesis can be derived from fixed tissues, where cells can only be analyzed from static images. The optical inaccessibility of the mammary fat pad has hampered efforts to image mammary epithelial cell behaviors at post-natal stages of development. Despite recent advances in deep tissue imaging (Dawson et al. 2021, Dawson & Visvader 2021), quantitative analyses of dynamic cellular behaviors driving branching morphogenesis have not been reported *in vivo* or in explant culture. Much of our mechanistic understanding comes from analysis of branch elongation in stroma-free organoid cultures. The tips of these branches are proliferative and multi-layered but lack the full myoepithelial coverage of TEBs in vivo (Ewald et al. 2008). Tracking epithelial cells in organoids has revealed that they engage in directional collective migration, but without breaching the basement membrane barrier (Ewald et al. 2012, Huebner et al. 2016). It has also been proposed that cell migration is coupled to radial cell intercalation to the surface epithelial layer to facilitate branch elongation (Neumann et al. 2018). Invasion of the tip is further propelled by mechanical constraint generated by myoepithelial cells behind the tip (Neumann et al. 2018). How these cell behaviors might change upon tip bifurcation, has not been studied in organoids. Furthermore, it is not known to what extent tip bifurcation is recapitulated in stroma-free culture conditions.

The process of tip bifurcation is perhaps best understood based on live imaging studies on ex vivo cultured embryonic organs (Goodwin & Nelson 2020, Lang et al. 2021). In the bifurcating kidney, differential morphogen receptor activation drives cell sorting movements between the branch point and daughter tips (Riccio et al. 2016). In developing lungs, proliferation is uniform in bifurcating tips, but oriented cell divisions and Myosin II (NMII) mediated contractility of the actin cytoskeleton may contribute to lung bifurcations (Schnatwinkel & Niswander 2013). In addition, differentiation of mesenchyme-derived smooth muscle cells at the leading edge of the tip was suggested to create a cleft between daughter tips (Kim et al. 2015). However, genetic inhibition of smooth muscle differentiation does not abrogate lung branching morphogenesis (Young et al. 2020). In the stratified tip of the salivary gland, localized deposition of fibronectin into a wedge-like structure opens a narrow cleft between cells to split the tip (Sakai et al. 2003, Larsen et al. 2006). More recent studies suggest that branching is driven by expansion and mechanical buckling of the surface cell sheet resulting from the combination of strong cell-matrix adhesion and weak basal cell-cell adhesions (Wang et al. 2021).

Here, we employ long-term high-resolution live imaging to investigate the cellular mechanisms driving expansion of the mammary epithelial network through terminal branch elongation and tip bifurcation. To discern cell behaviors during these processes, we took advantage of an ex vivo culture system that preserves the native tissue architecture with its epithelial-mesenchymal interactions (Kratochwil 1969, Voutilainen et al. 2012). Branching morphogenesis in this setup is consistent with morphometric analysis of embryonic mammary glands and includes both terminal bifurcation and side-branching events (Lindström et al. 2022). For a comprehensive view and mechanistic understanding of cell behaviors contributing to epithelial branching and elongation, we utilized a non-lineage specific approach to indiscriminately label and image the majority of embryonic mammary epithelial cells. Our findings suggest that terminal branching morphogenesis is characterized by coordinated changes in cell cycle activity and cell motility within the tip, to alternate between terminal branch elongation and branch point generation.

## Results

### Proliferation is localized to the central/luminal compartment and enriched in terminal tips

To investigate if localized cell proliferation contributes to embryonic mammary gland branching morphogenesis, we examined cell cycle status in fixed whole-mount glands at E18.5. We utilized a constitutive bicistronic fluorescent cell cycle reporter Fucci2a (Mort et al. 2014) and quantified cells in different phases of the cell cycle from 3D confocal images (**Figure 1A**). Cells preparing for mitosis (S/G2/M phase) express nuclear green fluorescent hGeminin-mVenus, while cells that exit mitosis (G1/G0 phase) express nuclear red fluorescent hCdt1-mCherry. Cells marked with either fluorophore constitute 75 ± 8 percent of all cells, as determined by manual counting of the cells from terminal segments of five different glands. For spatial analysis of whole glands, quantification was automated and the cell cycle status was expressed as the percentage of S/G2/M cells at different locations (**Figure S1A**). We first examined cell cycle activity based on cell distance to the surface of the epithelium. The number of cells peaked at a 4-5µm distance from the surface, as measured from the center of their nuclei, representing the position of the outermost cell layer (**Figure S1B**). Therefore, we considered cells with at ≤ 6µm and >6µm from the surface as basal and luminal cells, respectively. Surprisingly, we found that basal cells had a much higher G1/G0 cell cycle status overall compared to the luminal cells (**Figure 1B, Figure S1B**).

**Figure 1.**
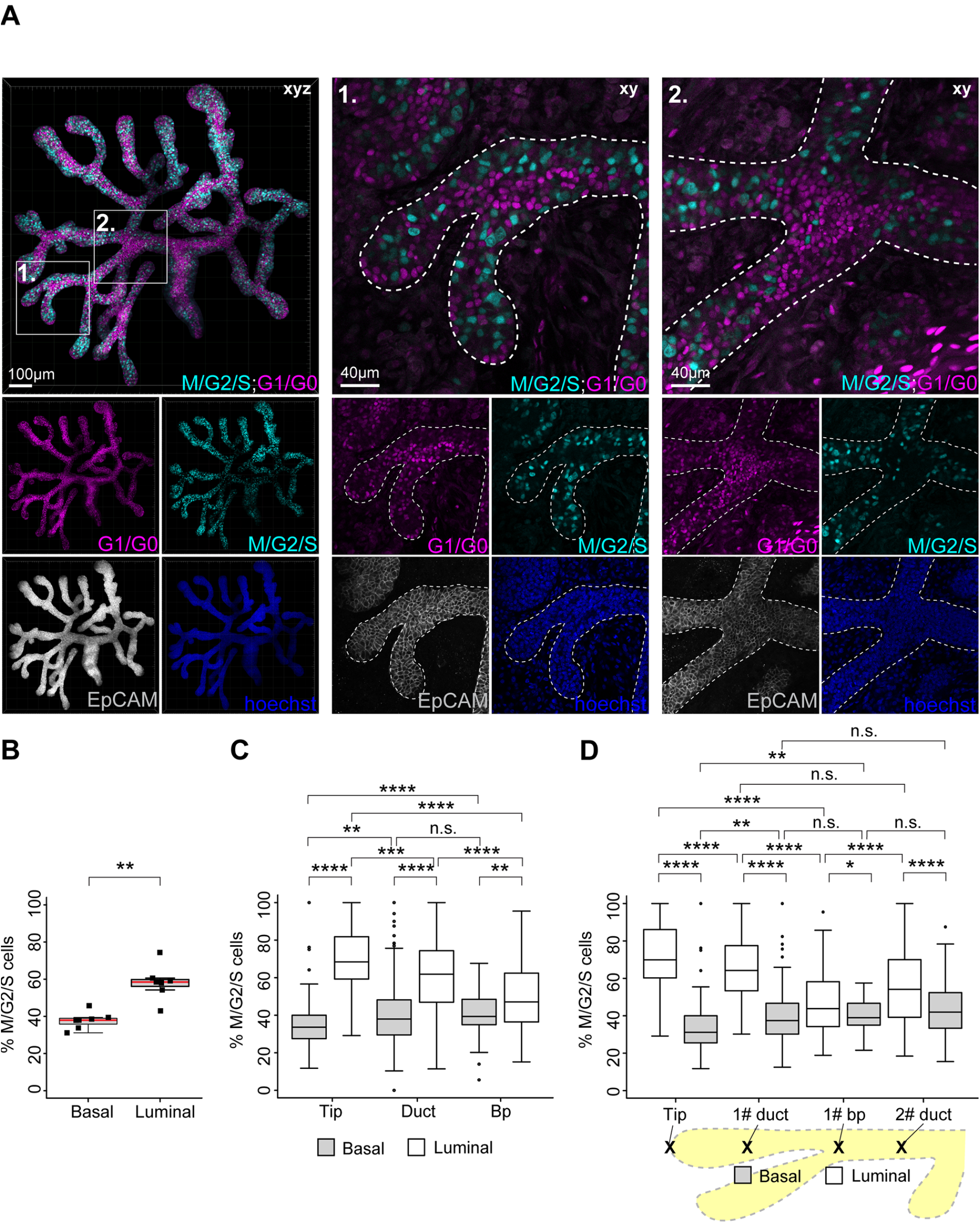
Proliferation is localized to the central/luminal compartment and enriched in terminal tips. **A)** Maximum intensity projection of the E18.5 mammary epithelium (left) with nuclear Fucci2a expression (M/G2/S; cyan, G1/G0; magenta) and EpCAM-staining (gray) used for epithelial surface rendering and masking of the mesenchyme. Cross-sections of two regions of interest are magnified on the right with both epithelial and mesenchymal signal visible: two terminal branches with their connecting branch point (1.) and a proximal branch point (2.). **B)** Percentage of cells expressing the green M/G2/S cell phase reporter of all Fucci2a-labeled cells in the “basal” outermost cell layer (≤ 6µm from the surface of the gland) and “luminal” inner epithelial compartment (> 6µm from the surface of the gland) (n = 7 glands). **C)** Percentage of cells in the M/G2/S phase in luminal and basal compartments of terminal tips (at ≤ 100µm distance from the terminal edge), ducts (within ≥ 50µm radius of the estimated center point) and branch points (within ≥ 50µm radius of the estimated center point) (n^Tips^ = 121, n^Ducts^ = 216, n^Branch points^ = 94, from 7 glands). **D)** Percentage of cells in the M/G2/S phase in the basal and luminal compartments of terminal branches (≥ 150µm in length, n = 74), with tips, ducts, branch points and the segment of duct behind the branch point separately analyzed. Data shown represent the median (line) with 25th and 75th percentiles (hinges) plus 1.5× interquartile ranges (whiskers). Statistical significance was assessed with the paired t-test (B), Wilcoxon’s signed rank test (C) and pairwise Wilcoxon’s signed rank test (D); *, P ≤ 0.05; **, P ≤ 0.01; ***, P ≤ 0.001; ****, P ≤ 0.0001.

Next, we assessed whether the cell cycle status was associated with any morphological domains of the mammary gland, and analyzed tips (the region within ≤ 100µm from the approximated terminal branch end point), ducts (the region within ≥ 50µm of the duct midpoint) and branch points (the region within ≥ 50µm radius of the center of the branch point) separately **(Figure 1A**; see close-ups 1-2, **Figure S1A**). Luminal cells displayed highest cell cycle activity in the tips, followed by ducts, while branch points were the least active (**Figure 1C**). Conversely, cell cycle activity of basal cells was lowest at the tip and lower than luminal cells in all the domains. As the network of ducts close to the tip are newly made, we explored whether cell cycle status changes with the distance to the closest tip. In both ducts and branch points, the proportion of S/G2/M luminal cells gradually decreased the further away they were from the tips while the basal cells displayed high G1/G0 status regardless of their location (**Figure S1C-D**). To investigate if differences in cell cycle status were established already during branch elongation and branching events, we focused our analysis on terminal branches only (**Figure 1A**, see close-up 1). The proportion of S/G2/M luminal cells peaked within a 100µm distance from the leading edge of the tip, while basal cells were lower in mitotic activity throughout the length of the branch (**Figure S1E**). Tips had a significantly higher percentage of S/G2/M luminal cells compared to the trailing duct and the latest branch point (**Figure 1D, Figure S1E**). Notably, branch points were even less mitotic than the segment of duct immediately behind the branch point. Taken together, cell cycle analysis of late embryonic mammary glands revealed that luminal cells are distally proliferative, while basal cells are relatively less-proliferative throughout the gland. Our data also suggest that luminal cells in the tips can be discerned from the latest branch point – a potential bifurcation site – by a much higher proliferation rate.

### Tips proliferate during branch elongation, while branch points become cell cycle repressed upon tip bifurcation

To examine bifurcating tips in detail, we identified tips where a cleft could be unequivocally detected in our dataset of fixed E18.5 mammary glands (**Figure 2A**). We first asked whether branches with bifurcating tips have a characteristic tip width or branch length, compared to branches with non-bifurcating tips, i.e. potentially elongating branches. Non-bifurcating tips were wider the longer the branch (**Figure 2B**), suggesting that tips widen as branches elongate in vivo. The width of the bifurcating tips was greater than non-bifurcating tips, irrespective of branch length, suggesting that tip widening is associated with tip bifurcation. Bifurcating tips were found in branches of all length, in line with the common notion that branching of the mammary gland is non-stereotyped (Goodwin & Nelson 2020, Scheele et al. 2017). To investigate if tip bifurcation involves local changes in cell cycle status, we quantified the Fucci2a cell cycle reporters in bifurcating tips, comparing the nascent daughter tips to the branch point in between (**Figure 2C**). We found that luminal cells in the branch point had a significantly reduced S/G2/M status compared to the daughter tips (**Figure 2D**), similar to branch points in terminal branches (**Figure 1D**). In contrast, the cell cycle status of the basal cells did not differ.

**Figure 2.**
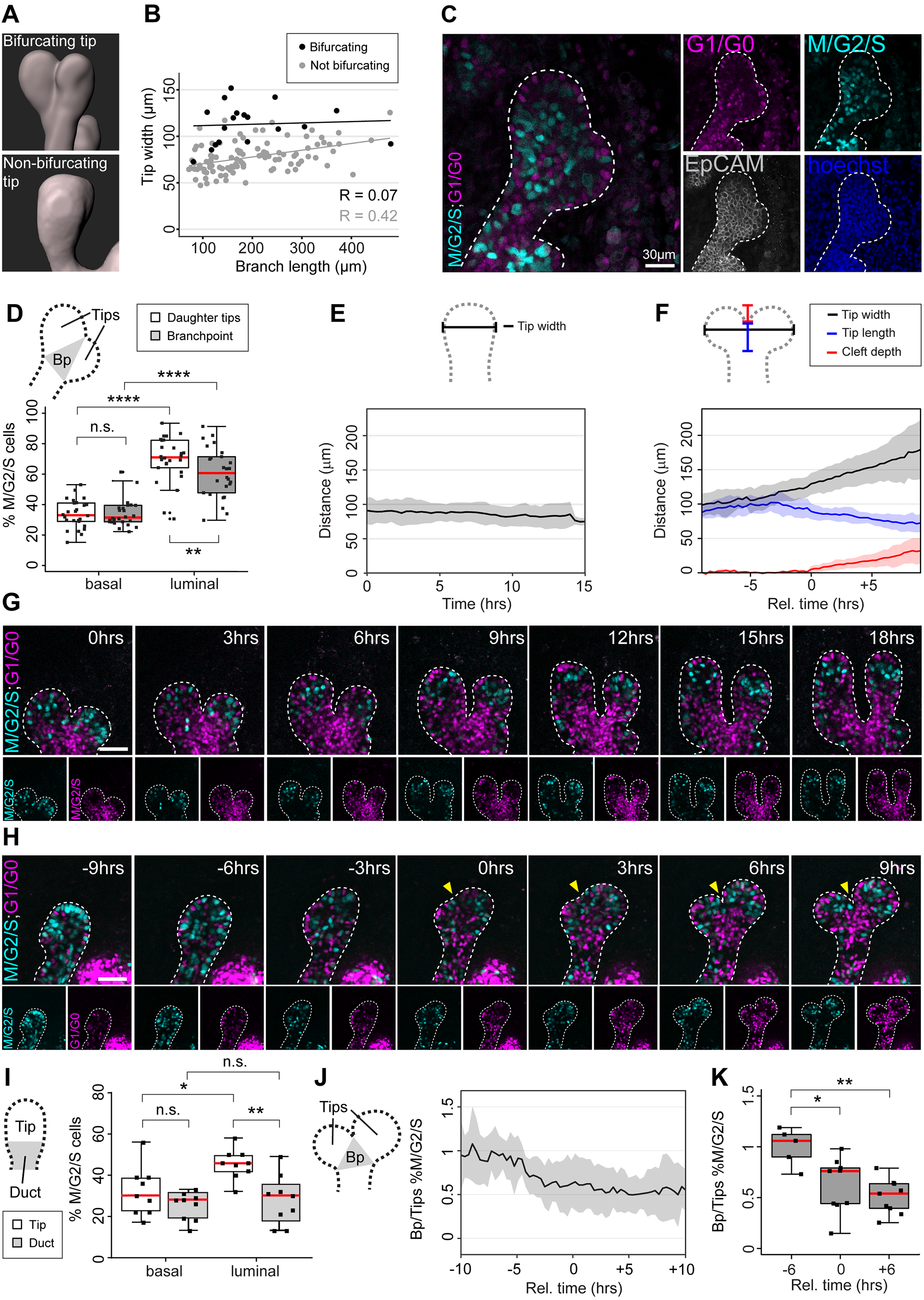
Tips proliferate during branch elongation, while branch points become cell cycle repressed upon tip bifurcation. **A)** Epithelial surface rendering of a bifurcating tip (top) and a non-bifurcating tip (below) generated based on whole-mount EpCAM staining. **B)** Analysis of tip width by branch length in terminal branches of the surface rendered E18.5 mammary gland with bifurcating and non-bifurcating tips. **C)** Confocal slice across a bifurcating tip and surrounding mesenchyme with Fucci2a expression (M/G2/S; cyan, G1/G0; magenta) and EpCAM-staining (grey). **D)** Quantification of cell cycle status in the “basal” and “luminal” compartment of bifurcating tips based Fucci2a fluorescent nuclei, comparing the emerging daughter tips to the branch point as delineated in the schematic (n = 26). **E)** Confocal time-lapse imaging was performed on ex vivo cultured mammary glands with epithelial Fucci2a expression (K14-Cre; Fucci2a), and tip width was measured as shown in the schematic on top from elongating terminal branches, based on an epithelial surface rendering (n = 9). **F)** Dimensions of the bifurcating tip were measured as indicated in the schematic from time-lapse videos at different time points before and after the appearance of a cleft (T0) (n = 10). **G)** Captions of a time-lapse imaging series featuring two elongating branches with epithelial Fucci2a expression. Note the enrichment of M/G2/S phase nuclei in the tips, which was used as a proxy to define the tip border in the absence of a discernable neck. **H)** Captions featuring a bifurcating tip with epithelial Fucci2a expression (yellow arrowhead points to the cleft). **I)** Quantification of cell cycle status (% M/G2/S phase nuclei) from confocal time-lapse imaging videos of elongating branches comparing the tip to the trailing duct with “basal” and “luminal” compartments analyzed separately (n = 9, data was pooled from all time points). **J)** The relative cell cycle status between the branch point and the daughter tips (the branch point-to-tip ratio of % M/G2/S nuclei) of time-lapse imaged bifurcations was plotted against time normalized according to the appearance of the cleft (T0) (n = 9). **K)** The ratio of % M/G2/S nuclei was compared 6 hours before, upon (T0) and 6 hours after the appearance of the cleft in different videos (n = 9). Data shown in D, I and K represent the median (line) with 25th and 75th percentiles (hinges) plus 1.5× interquartile ranges (whiskers). Data shown in E, F and J represent average (line) ± S.D (shaded region). Statistical significance was assessed with the pairwise Wilcoxon’s signed rank test (D) and paired t-test (I & K); *, P ≤ 0.05; **, P ≤ 0.01; ***, P ≤ 0.001; ****, P ≤ 0.0001.

To study what happens between branch elongation and tip bifurcation in real time, we performed live imaging of *ex vivo* cultured mammary glands derived from embryos expressing the Fucci2a reporter only in epithelial cells (K14-Cre;R26R-Fucci2-flox/flox mice). We utilized an established protocol, where E13.5 mammary buds isolated with their surrounding mesenchyme, initiate branching morphogenesis on day 3 of culture (**Figure S2A**; Voutilainen et al., 2012; Lan et al. 2022). Confocal whole-mount imaging was performed on day 5-6, when branching events are frequent using a set-up where the explants were attached to a transparent air-permeable membrane at the bottom of a dish and covered in media (**Figure S2B**). Cells were monitored and analyzed in elongating branches and in branches undergoing terminal bifurcation for up to 28 hours at 20 min intervals (**Figure S2C-E, Videos 1-2**). We first studied the morphometrics of the tip based on epithelial 3D surface rendering, created around all the fluorescent nuclei. Tips of elongating branches that did not bifurcate during the time-lapse maintained a constant width over time (**Figure 2E**). Tips that bifurcated increased in width before a cleft became visible (**Figure 2F**). At the same time, the neck of the tip became more pronounced, and the tip flattened as measured from the neck to the leading front.

To study cell cycle dynamics, Fucci2a fluorescent nuclei were automatically detected as spots and quantified. Consistent with our in vivo findings, tips of elongating branches appeared to be populated with S/G2/M phase luminal cells (**Figure 2G, Video 1**), which we used as a proxy to distinguish the tip from the more G1/G0 positive trailing duct. We noticed that prolonged imaging had a negative effect on cell cycle activity over time, but the difference between the tip and the trailing duct persisted throughout the imaging (**Figure S2F**). The aggregated time-lapse data from all the videos confirmed the enrichment of proliferative luminal cells in the tip, consistent with our in vivo findings (**Figure 2I**). To assess if the cell cycle status changes within the tip as tips bifurcate (**Figure 2H, Video 2**), we determined the proportion of S/G2/M cells in branch points and daughter tips and plotted their ratio as a function of time, where the time point of cleft appearance was defined as T0 (**Figure 2J**). The branch point/tip ratio decreased substantially prior to appearance of the cleft and remained low for several hours as bifurcation advanced (**Figure 2J-K**). These results point towards a localized cell cycle repression at the branch point. However, the cell cycle status might also change because cells move in and out of the branch point in cell cycle dependent manner. To investigate this possibility, we compared the proportion of incoming and exiting S/G2/M and G1/G0 cells and found no difference (**Figure S2G-H**).

Taken together, our real time cell cycle analysis of ex vivo cultured embryonic mammary glands indicates that elongating branches maintain a pool of S/G2/M luminal cells at the tip. As the tip bifurcates, cells at the branch point become cell cycle repressed, while the newly formed daughter tips remain proliferative.

### Inhibition of cell proliferation does not prevent branch elongation or branch point generation

Cell proliferation is required for growth of the mammary epithelium, but whether cell divisions *per se* can elongate or initiate branches during branching morphogenesis, remains unclear. To address this question, we examined the effects of global cell cycle inhibition on branching morphogenesis by utilizing a short 2-hour ex vivo treatment of Mitomycin C (MMC) followed by culture in fresh medium for 24 hours using explants expressing epithelial GFP (K14-Cre;mT/mG) for easy visualization (**Figure 3A**). EdU labeling at the end of the culture period confirmed that cell proliferation had been suppressed (**Figure 3B**). Next, we quantified the number of new branches generated over the 24-hour culture after MMC treatment. As expected, the number of new branches was much lower than in non-treated control explants (**Figure 3C**).

**Figure 3.**
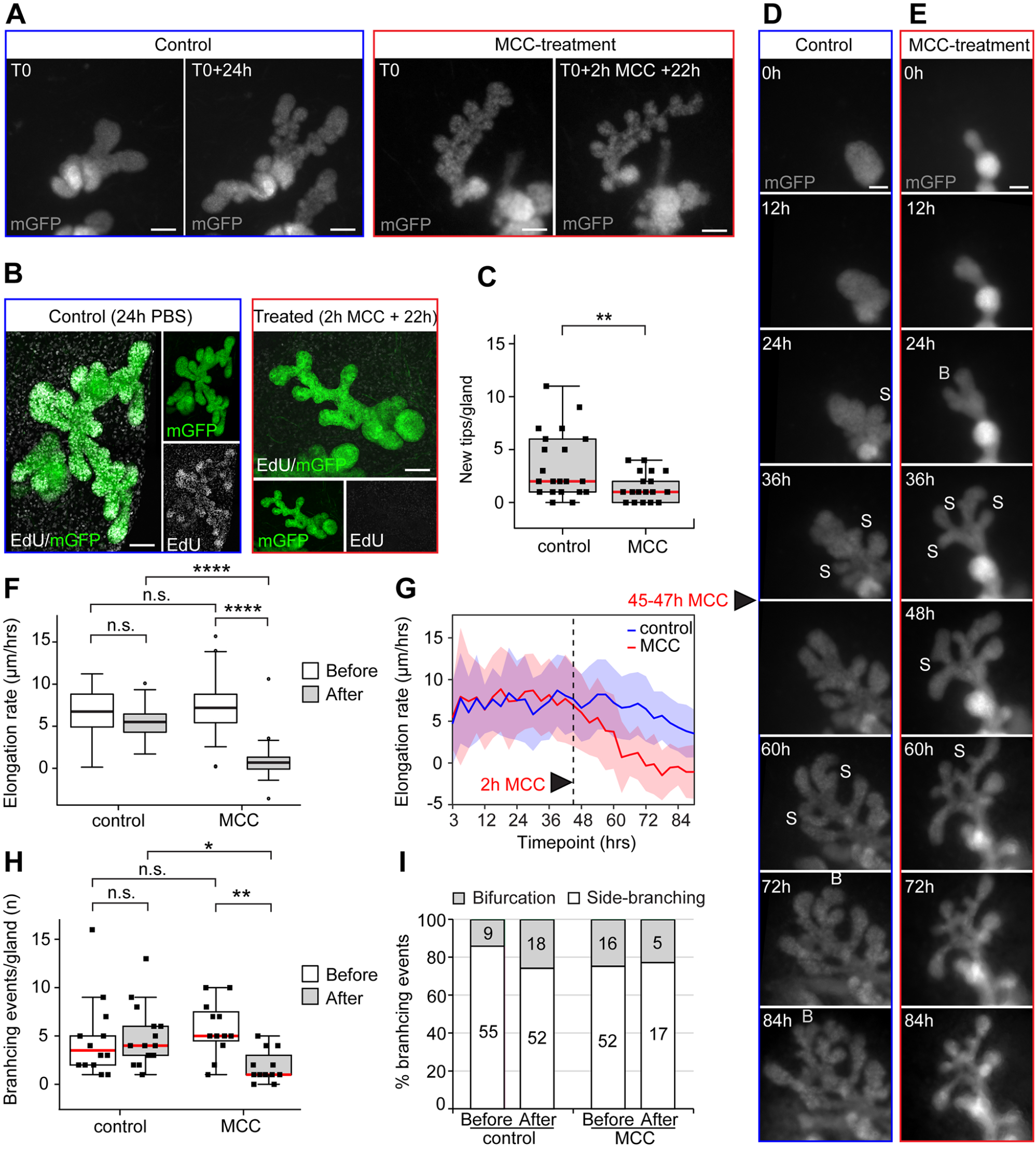
Inhibition of cell proliferation does not prevent branch elongation or branch point generation. **A)** Ex vivo cultured mammary glands with epithelial mGFP expression (K14-Cre;R26R-mTmG), were imaged before and after 2 hour treatment with 1µg/ml of Mitomycin C (MCC) for cell cycle repression or vehicle (control) followed by 22h of culture in normal media (mGFP = grey). **B)** To assess the effects of MCC-treatment on cell proliferation, control and treated glands were labeled with EdU after the culture period, and fixed for staining and confocal imaging (maximum intensity projections of EdU in white and mGFP in green). **C)** The number of tips was quantified from wide-field images of the mGFP-labeled epithelium (A) before and after culturing (n^control^ = 21 glands from 10 explants, n^treated^ = 22 glands from 12 explants; four experiments). The effects of cell cycle repression on branching morphogenesis was studied by wide-field time-lapse imaging of epithelial mGFP expressing explants (93 hours of imaging with 3-hour intervals). In the middle of the time-lapse (45-47 hours), explants were treated for two hours either with vehicle **(D)** or with MCC **(E)**, followed by imaging in normal media. **F)** The length of terminal branches was measured as a function of time and the elongation rate (Δ length/time) between control and treated glands, before and after treatment compared (n^control^ = 73 branches from 13 glands, n^treated^ = 62 branches from 12 glands; two time-lapse experiments). **G)** The rate of elongation between adjacent time points was determined for each branch and the average plotted as a function of time. **H)** Branching events were scored in each gland by careful observation of time-lapse videos and the number of events that occurred in control and treated glands before and after treatment compared (n^control^ = 13 glands, n^treated^ = 12 glands). The branching events could be further identified as bifurcations or side-branching events as shown in D and E. **I)** To assess their prevalence, the overall percentages of different types of branching events were calculated by pooling of events from different glands (total number of events indicated in the chart). Data shown in C, F and H represent the median (line) with 25th and 75th percentiles (hinges) plus 1.5× interquartile ranges (whiskers). Data shown G represents average (line) ± S.D (shaded region). Statistical significance was assessed with the Wilcoxon’s signed rank test (C), paired t-test (F) or paired Wilcoxon’s signed rank test (H); *, P ≤ 0.05; **, P ≤ 0.01; ****, P ≤ 0.0001. Size bars = 100 µm.

To analyze where and when the effects of MMC were manifested, we performed low-resolution time-lapse imaging of cultured mammary explants for a total of 93 hours, taking pictures every 3 hours (**Figure 3D-E, Video 3-4**). A two-hour Mitomycin C treatment was administered to half of the explants at 45 hours, when several branches had already formed. We first analyzed elongation of branches by measuring branch length over time. The rate of elongation was significantly reduced in MCC-treated samples compared to control samples (**Figure 3F**). More detailed analysis revealed that branch elongation did not cease completely until 16 hours after the treatment (**Figure 3G**). This finding prompted us to analyze the role of cell divisions in initiation of branch formation. As tip bifurcation and side-branching are essentially different forms of tissue morphogenesis (Wang et al. 2017), we wanted to investigate how cell divisions contribute into each. As expected, time-lapse imaging revealed that MMC treatment reduced branching events (**Figure 3H**), yet the ratio of tip bifurcation and side-branching events remained unchanged (**Figure 3I**), indicating a similar response to the treatment. Most of the remaining branching events occurred within 16 hours of the MMC-treatment (**Figure S4A-B**), suggesting that branching beyond this point is limited by exhaustion of new cells. Interestingly, we did notice a difference in the stability of branches generated via tip bifurcation and side-branching after the treatment. Side-branches were stable throughout imaging, while tip bifurcations often returned to their un-bifurcated state (**Figure 3J**).

Based on imaging the effects of cell cycle repression in ex vivo culture, we found no support for the role of cell divisions in driving branch elongation or triggering branching events *per se*. However, branching morphogenesis is progressively limited by cell number. In particular, cell proliferation is required to sustain two daughter tips after a tip bifurcation has taken place.

### Epithelial cell contractility is essential for mammary gland branching morphogenesis

An important regulator of epithelial shaping is the actomyosin network, where NMII generates forces by crosslinking and pulling on actin filaments (Miao & Blankenship 2020). To investigate its role in the morphogenesis of bifurcating tips, we performed whole mount phalloidin staining and focused on regions of high epithelial curvature, where actin bundles might be enriched, and cells engaged in contractile activity (**Figure 4A-B**). Epithelial and mesenchymal intensity was analyzed separately, based on rendering of the epithelium. Quantification of F-actin intensity in three gross areas: around the cleft, daughter tips and neck (**Figure 4C**) revealed no difference, although staining was slightly elevated in the epithelium compared to the mesenchyme. These quantifications were in line our observation that tips in vivo, varying from early to later stages of bifurcation (**Figure 4A-B**), displayed no signs of localized F-actin-mediated tissue constrictions around the cleft or neck area. However, these findings do not exclude the possibility that dynamic changes in actomyosin contractility could be important at cellular and sub-cellular level. To functionally assess the role of actomyosin contractility, we inhibited NMII activity in ex vivo culture by treating mammary explants with blebbistatin over a 24-hour period and quantified the number of new branches in control and drug-treated samples (**Figure 4D-E**). The results show that branching morphogenesis was strongly inhibited by blebbistatin, although growth of the epithelium appeared not to be affected (Figure 4F). To distinguish whether contractility is required in the epithelium or mesenchyme, we inhibited NMII activity with blebbistatin in mesenchyme-free organoids using a novel technique to culture intact E16.5 mammary epithelial sprouts in Matrigel in stroma-free conditions (Lan et al. 2022). Organoids were cultured in Matrigel for 2 days, followed by a 24-hour treatment with blebbistatin (**Figure 4G**). To our surprise, blebbistatin strongly inhibited branching of epithelial organoids (**Figure 4H**). Again, epithelial growth was unobstructed as determined by measuring the epithelial area and based on EdU staining **(Figure 4I-J**). These results indicate that mammary gland branching morphogenesis is dependent on epithelial actomyosin contractility.

**Figure 4.**
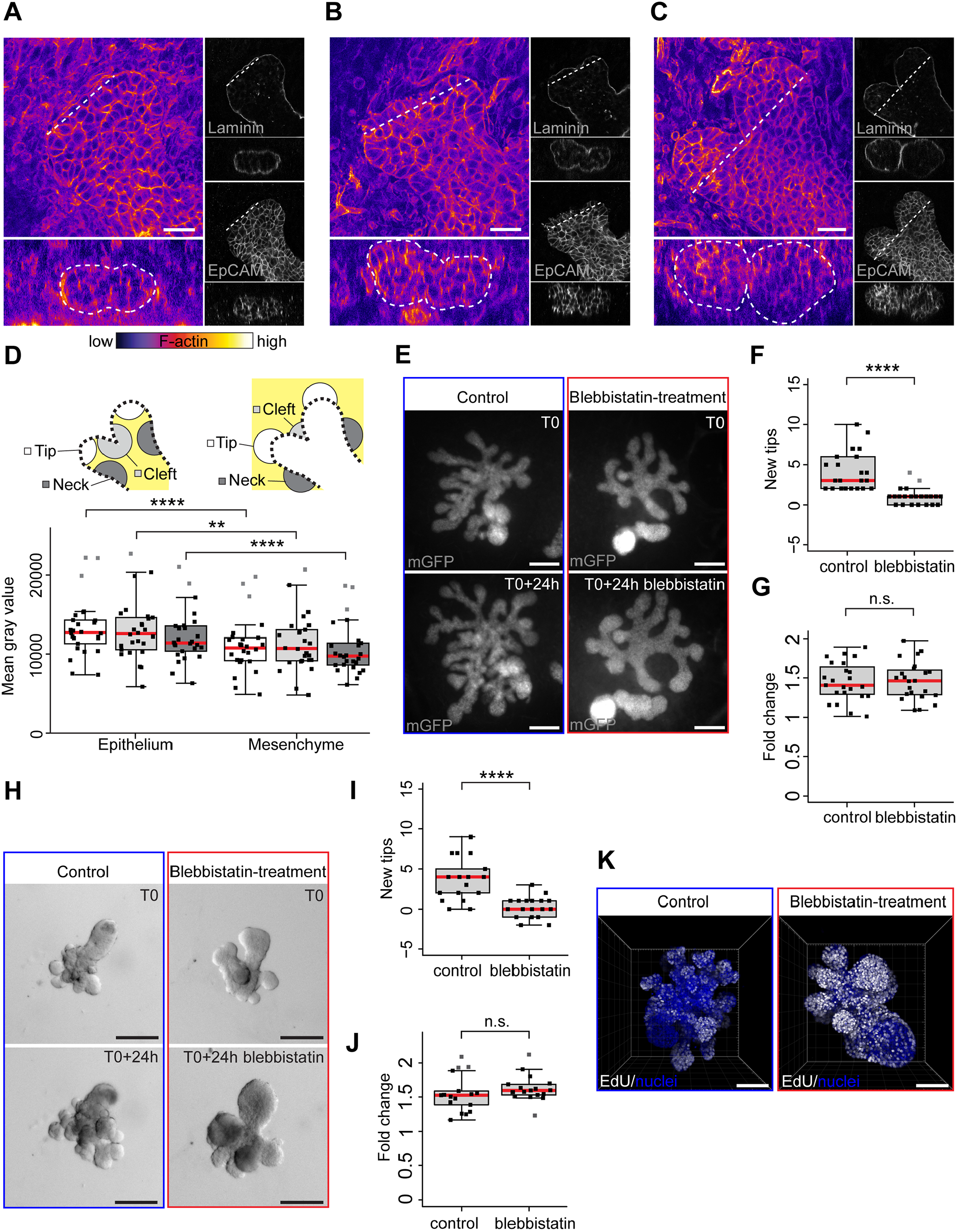
Epithelial cell contractility is essential for mammary gland branching morphogenesis. **A)** Confocal xy-slice of a tip at an early stage of bifurcation, stained with F-actin- (Fire LUT), laminin (grey) and EpCAM (grey). A vertical cross-section of the tip along the dotted line in the xy-view is shown below. Similar images of an early bifurcating tip with a more obvious cleft (B) and late bifurcating tip, where cleft has progressed further (C). **D)** Quantification of F-actin intensity within epithelial and mesenchymal regions around the daughter tip, cleft and neck of the bifurcating tip as marked in the schematic (n = 26, from 16 glands of 7 embryos). **E)** Ex vivo cultured mammary glands with epithelial mGFP expression (K14-Cre; R26R-mTmG), were imaged before and treatment with 20µg/ml of blebbistatin or vehicle (mGFP = grey). The number of tips was quantified **(F)** and the growth area measured **(G)** from wide-field images of the mGFP-labeled epithelium before and after treatment (n^control^ = 22 glands from 10 explants, n^treated^ = 22 glands from 10 explants; two experiments). **H)** Epithelial organoids cultured in 3D Matrigel™ were imaged before and after treatment with 20µg/ml of blebbistatin or vehicle (brightfield). The number of tips was quantified **(I)** and the growth area measured **(J)** from brightfield images before and after treatment (n^control^ = 17 glands, n^treated^ = 18 glands, from two experiments). To assess the effects of blebbistatin-treatment on cell proliferation, control and treated organoids were labeled with EdU after the culture period, and fixed for staining and confocal imaging (maximum intensity projections of EdU in white and hoechst in blue). Data shown represents the median (line) with 25th and 75th percentiles (hinges) plus 1.5× interquartile ranges (whiskers). Statistical significance was assessed with the paired Wilcoxon’s signed rank test (D), Welch t-test (F, J) or Wilcoxon’s signed rank test (G, I); **, P ≤ 0.01; ****, P ≤ 0.0001. Size bars in A-C = 30 µm, in E & H = 200µm and in K = 100µm.

### Differential cell motility creates a retrograde flow of cells during branch elongation, which is restricted upon tip bifurcation

To address whether branch elongation involves epithelial cell motility, we surmised that mammary cells would have a motile phenotype, characterized by an elongated rather than cubical epithelial cell shape. To analyze cell shapes in 3D, mammary epithelial cells were sparsely labeled with cytosolic tdTomato using a doxycycline inducible K5-rtTA;TetO-Cre and analyzed in fixed glands. We observed that most cells had a spindle-like shape with pointed edges (**Figure 5A**). Cells at the tip, closer to the leading front of elongation (≤ 100µm), were less spherical than cells further away in the duct (**Figure 5B**). To study cell motility in elongating branches, we performed automated cell tracking based on nuclear Fucci2 expression in the live imaging dataset (**Video 5**). As the Fucci2a expression is temporarily lost during mitosis, we did not attempt to construct cell lineages by combining tracks based on different Fucci2a fluorophores. The length of the tracks generated was similar for both fluorophores, with a median track length of 3,7 hours for Fucci2a-green and 4 hours for Fucci2a-red cells (tracks shorter than two hours were excluded) (**Figure S4A**). Overall, cells in the M/G2/S phase of the cell cycle were slightly faster and more persistent compared to cells in G1/G0 phase of the cell cycle (**Figure S4B-D**). To assess if any differences exist in cells’ velocity along the elongating branches, we mapped all tracks based on their position with respect to the tip (**Figure 5C**). Cells in the tip, defined as the region with higher proportion of M/G2/S phase cells, were significantly faster and more persistent compared to the cells in the trailing duct (**Figure 5D, Figure S4E-F, Video 5**). The velocity of the tracks formed a descending gradient with increasing distance from the leading front of the tip (**Figure 5E**).

**Figure 5.**
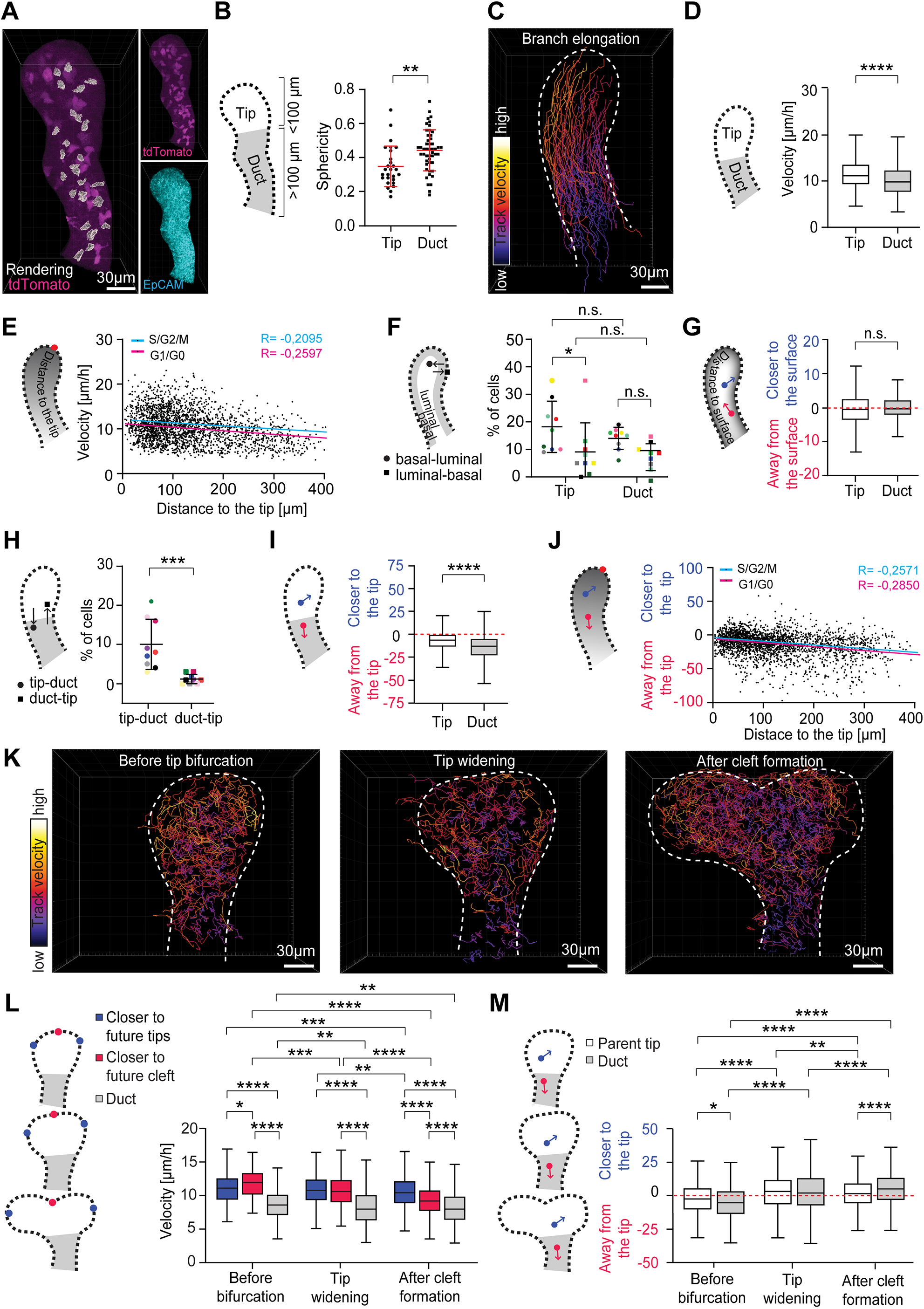
Differential cell motility creates a retrograde flow of cells during branch elongation, which is restricted upon tip bifurcation. **A)** Maximum intensity projection of sparsely labeled tdTomato+ cells (tdTomato-fl/fl, induced by Krt5-rtTA;TetO-Cre in the presence doxycycline) in the E18.5 mammary epithelium (magenta), together with volume rendering of isolated cells (gray). EpCAM staining (cyan, bottom right) was used for creating an epithelial surface rendering to mask mesenchymal and to highlight epithelial tdTomato expression (magenta, top right). **B)** Quantification of tdTomato+ cells sphericity in the tip (at ≤ 100 µm distance from the landmark point in the tip) and duct (at > 100 µm distance from the landmark point in the tip) (n^tip^ = 27 cells, n^duct^ = 49 cells, from 9 branches). **C)** Visualization of Fucci2a cell tracks, color-coded for mean velocity (fire LUT, black→white = low→high), in an elongating mammary epithelial branch from a 3D time-lapse confocal video of an ex vivo cultured mammary explant (K14-Cre;Fucci2a). **D)** Comparison of mean velocity of tracked cells between tip (defined as the region visibly enriched with M/G2/S cells) and duct (n^tip^ = 600 cells, n^duct^ =1814 cells, from 9 elongating branches). **E)** Distribution of mean velocities of cell tracks with respect to their distance to the leading edge of the tip, where the trends of M/G2/S and G1/G0 tracks are shown as cyan and magenta lines, respectively (n^M/G2/S^ =785, n^G1/G0^ =1629, from 9 branches). **F)** Percentages of cell tracks per video in the tip and duct that switch from “basal” outermost cell layer (≤ 6µm from the surface of the gland) to “luminal” inner epithelial compartment (> 6µm from the surface of the gland) and vice versa (n = 9 branches). **G)** Tendency of “luminal” cells in the tip and duct to move closer to the epithelial surface (calculated based on cell distance to the epithelial surface) in elongating branch (n^tip^ = 403 cells, n^duct^ = 1212 cells; from 9 branches). **H)** Percentages of cell tracks per video that start within the tip (M/G2/S-enriched region) and end in the duct and vice versa (n = 9 branches). **I)** We analyzed if cells at the end of their tracks were closer or further away from the leading edge of the tip than where they started (change in cell distance to the tip; positive values = closer, negative values further) and compared cells that started within the tip and duct (n^tip^ = 600 cells, n^duct^ =1814 cells, from 9 branches). **J)** The change in cell-leading edge distance was plotted against the starting position of the tracks with the trends of M/G2/S and G1/G0 tracks shown as cyan and magenta lines, respectively (n^M/G2/S^ =785 cells, n^G1/G0^ =1629, from 9 branches). **K)** Visualization of Fucci2a cell tracks, color-coded according to mean velocity (fire LUT), before tip bifurcation (left), during tip widening (middle) and after cleft formation (right) from 3D time-lapse confocal videos of ex vivo cultured mammary explants (fire LUT, black→white = low→high). **L)** Mean velocity of tracks in bifurcating tips, divided in two groups based on whether they were closer to future tips (blue) or future cleft (red) and compared to those in the duct (grey) according to whether they started before bifurcation (n^tip^ = 272, n^cleft^ = 77, n^duct^ = 154), during tip widening (n^tip^ = 688, n^cleft^ = 279, n^duct^ = 1152) or after the cleft became visible (n^tip^ = 1098, n^cleft^ = 1143, n^duct^ = 1645). **M)** We analyzed if cells at the end of their tracks were closer or further away from the leading edge/future cleft than where they started and compared cells that started within the parent tip and duct before bifurcation (n^tip^ = 346, n^duct^ = 174), during tip widening (n^tip^ = 879, n^duct^ = 1240) and after the cleft first appeared (n^tip^ = 1997, n^duct^ = 1534). Data shown in D, G, I, L & M represents the median (line) with 25th and 75th percentiles (hinges) plus 1.5× interquartile ranges (whiskers). Data shown in B, F and H represents average ± S.D. Statistical significance was assessed with the Student’s t-test, Mann-Whitney test (B-I) and Wilcoxon’s signed rank test (L-M); *, P ≤ 0.05; **, P ≤ 0.01; ***, P ≤ 0.001; ****, P ≤ 0.0001.

Next, we explored how differential cell motility affects cells’ contribution to different tissue domains. First, we investigated if cell movement contributes to epithelial surface expansion and analyzed the movement of cells between basal and luminal ‘compartments’. We observed a trend where the basal cells moved more often to the luminal compartment than vice versa, regardless of their position along the branch – tip or duct (**Figure 5F**). To assess if there was any overall trend of radial movement of luminal cells, we analyzed the position of cells relative to the epithelial surface and found that overall, their positions were maintained (**Figure 5G**). The gradient of cell velocities in the elongating branch (**Figure 5E**) indicated that slower tip cells may get left behind and end up in the trailing duct. To investigate this possibility, we analyzed the destination of the tip and duct cells and found that significantly more tip cells ended up in the duct than duct cells in the tip (**Figure 5H**). To evaluate cell lag along the elongating branch, we measured the distance of cells to the leading edge of the tip at the start and end of the track: cells in the trailing duct lagged behind significantly more than cells in the tip (**Figure 5I**). The further away from the leading edge the cell was the greater was also the lag (**Figure 5J**).

As the front/center part of the tip transforms into the branch point once the tip bifurcates, we hypothesized that cell movement may change at this location. Therefore, we examined cells’ velocity in branches undergoing terminal bifurcation (**Figure 5K, Video 6**). Remarkably, once the cleft had been established, cells at the branch point appeared to slow down. To confirm this observation, we analyzed tracks in the parent tip based on their proximity to landmarks positioned at the future daughter tips and the emerging cleft and stratified the tracks based on when they were generated – during branch elongation, tip widening, or cleft formation. Tracks that were closer to the future cleft became slower, while the cells closer to the daughter tips remained motile (**Figure 5L, Figure S4G-J**). Once the cleft had been established, the branch point cells were significantly slower than the cells of the daughter tips, suggestive of their localized immobilization. We hypothesized that immobilization of the cells at the branch point may restrict the flow of lagging cells to the trailing duct, preventing its further elongation. Indeed, we found that the flow of cells from the tip towards the trailing duct was significantly reduced during tip bifurcation (**Figure 5M**).

In conclusion, analysis of cell behaviors in elongating branches revealed a gradient of cell movement along the elongating branch. This creates a retrograde flow of cells that contributes to the elongation of the trailing duct. Once a branch point is generated, the flow of the cells is interrupted, possibly due to immobilization of the branch point cells.

### The invasion of the tip is facilitated by directional cell movement, which is redirected during tip bifurcation

Branch elongation in the mammary gland involves the forward advancement and invasion of the tip into the surrounding stromal tissue, which defines the direction of the branch. The velocity of tip cells correlated with the velocity of the tip itself (**Figure 6A**), implying that tip advancement may be propelled by cell movement. To investigate further if the direction of cell movement is consistent with the direction of tip advancement, we analyzed the vectors of cell displacement in the elongating branch (**Figure 6B**). We measured the angle of the cell displacement vectors to the vector of tip displacement and found that bulk of the cells moved to the same direction as the tip (**Figure 6C-D**). Furthermore, tip cells displayed significantly smaller angles than the cells in the trailing duct (**Figure 6C-D**), suggesting that their movement was more aligned with the direction of the tip. These results indicate that the direction of the invading tip may be governed by directionality of the tip cells.

**Figure 6.**
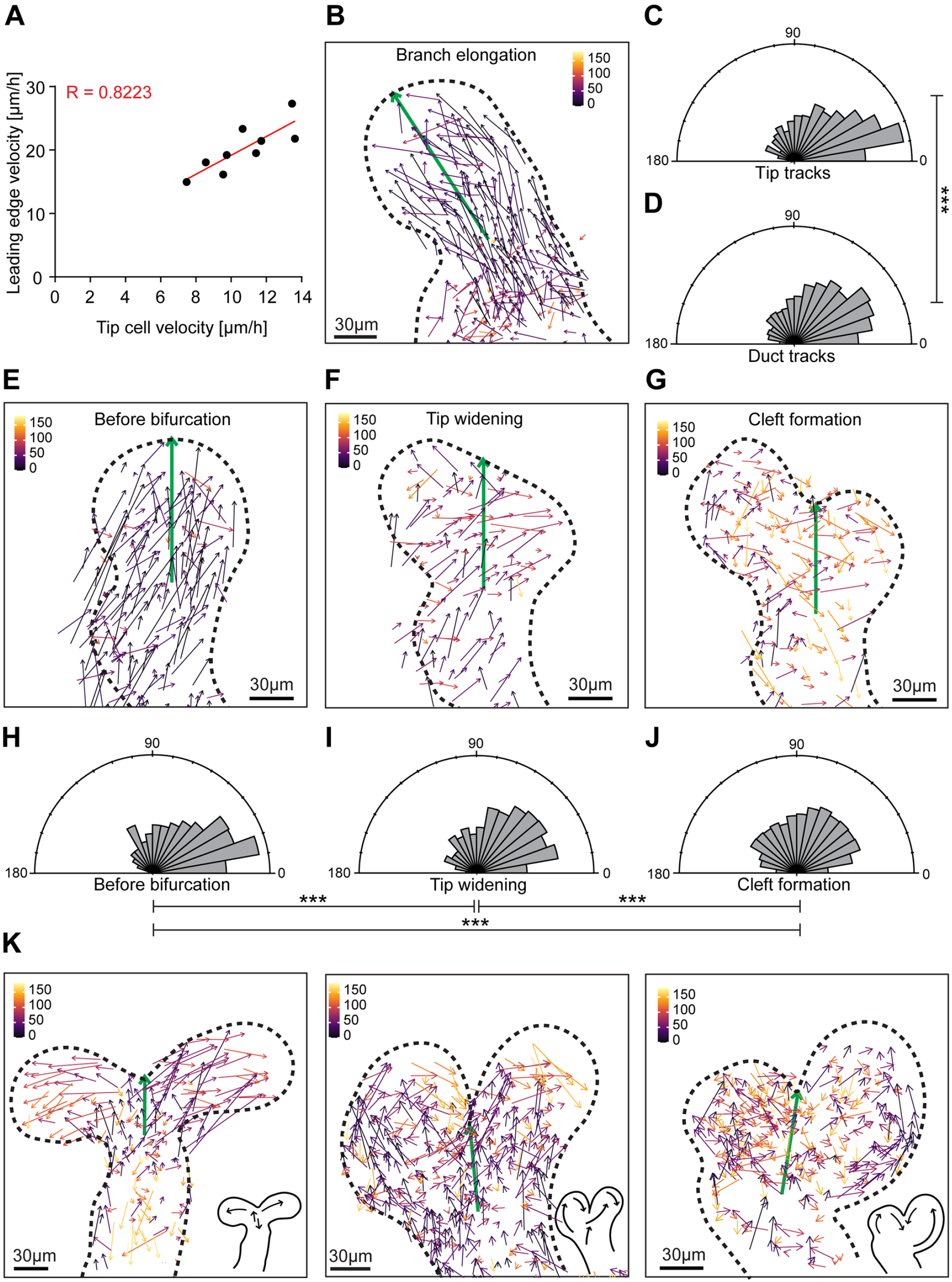
The invasion of the tip is facilitated by directional cell movement, which is redirected during tip bifurcation. **A)** Correlation of leading edge velocities of elongating branches with the average velocity of tip cell tracks (n = 9). **B)** Cell displacement vectors, color-coded based on their angle (fire LUT, black = 0°, white = 180°) to the displacement vector of the leading edge the elongating branch (green). Low angles indicate parallel direction of movement, while high angles reflect anti-parallel direction of movement. Rose diagram of the angles of tip cell displacement vectors **(C)** and duct cell displacement vectors **(D)** against the displacement vector of the leading edge as a reference (n^tip^ = 600, n^duct^ = 1814). Cell displacement vectors that were created before bifurcation **(E)**, during tip widening **(F)** and after cleft formation **(G)**, color-coded based on their angle (fire LUT, black = 0°, white = 180°) to the reference vector (green) running from the center of the neck to the leading edge/future cleft. Rose diagram of the angles of tip cell displacement vectors relative to the reference vector, depending on whether they started and ended before bifurcation (n = 214) **(H)**, during tip widening (n = 475) **(I)** and after the cleft first became visible (n = 2214) **(J). K)** Examples of cell displacement vectors from different bifurcations during cleft formation, showing variable patterns of directional cell movement. Groups of cell displacement vectors with a similar direction (similar angles against the green reference vector) were indicated as “cell streams” relative to the outline of the tip on the lower right corner. Uniformity of angles was tested with the Rayleigh test and statistical significance between groups was assessed with the Watson’s Two-Sample Test of Homogeneity; *, P ≤ 0.05; **, P ≤ 0.01; ***, P ≤ 0.001; ****, P ≤ 0.0001.

Upon tip bifurcation, two daughter tips are generated that begin to elongate branches in two different directions. To explore how this is reflected at the cellular level, we visualized cell displacement vectors in bifurcating tips (**Figure 6E-G**). We observed that the direction of the cells gradually became more dispersed, which correlated with the emergence of the daughter tips. We compared the displacement vectors of the cells to a reference vector, running from the center of the neck to the future cleft, and calculated the angle in between. We found that before widening of the tip, cells were moving parallel to reference vector (**Figure 6H**), much like in the elongating branches (**Figure 6B-D**). However, once the tip begun to widen and the cleft became visible, the angles became significantly wider, indicating that cells moved more perpendicular to the parental tip axis (**Figure 6I-J**). Examining cell displacement vectors in different bifurcating tips revealed two separate streams of perpendicular cell movement entering into the daughter tips (**Figure 6K**). Interestingly, anti-parallel cell movement was often (but not in every case) observed at one or both sides of the cleft.

In conclusion, these results suggest that invasion of the tip is mediated by directional cell migration, which becomes redirected between the two daughter tips during tip bifurcation.

### Tip bifurcation involves narrowing of the cleft and is independent of the mesenchyme

Tip bifurcation involves establishment of a cleft between the two daughter tips that becomes a permanent part of the branch point. In other organs, the cleft has been described either as a localized obstruction or a wedge that splits the tip (Sakai et al. 2003, Kim et al. 2015). To visualize clefts in vivo, we performed whole mount 3D imaging of the basement membrane visualized with the laminin antibody (**Figure 7A**). Measurements of the angle and depth of the cleft showed that shallow (i.e. nascent) clefts were associated with wider angles and deeper clefts with narrower angles (**Figure 7B**). To monitor clefting in real time, we used epithelial cell surface rendering to measure cleft angles in bifurcating tips and found that the cleft was initially wide and became narrower over time (**Figure 7C-D**), suggesting that clefting in the mammary gland is a process of gradual narrowing. To investigate to what extent cleft formation contributes to tip bifurcation, we analyzed movement of the cleft and daughter tips. Visualization of the epithelial outline revealed that both advancement of the daughter tips and ingression of the cleft contributed to the shaping of the tip (**Figure 7E-F**). The outward movement of the daughter tips and the inward movement of the cleft was almost equal, suggestive of their mutual contribution (**Figure 7F**).

**Figure 7.**
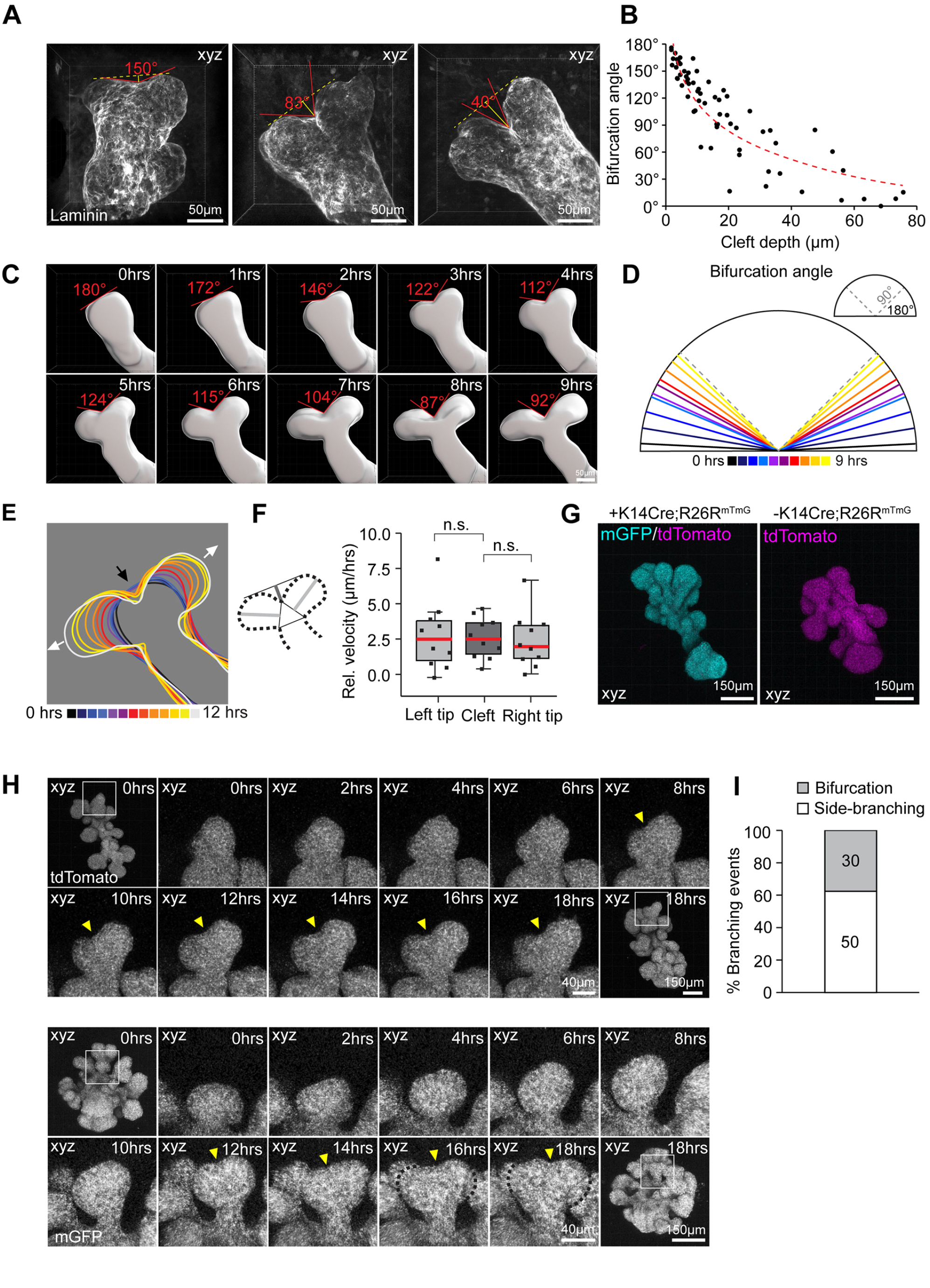
Tip bifurcation involves narrowing of the cleft and is independent of the mesenchyme. **A)** Maximum intensity projections of bifurcating tips, imaged from E17.5 – E18.5 mammary glands, with a shallow cleft (left) and deeper clefts (middle, right) visible based on laminin-staining. **B)** The angle and depth of the cleft was measured as indicated in (A) and plotted for 65 bifurcations from 30 glands of 13 embryos. **C)** Images of an epithelial surface rendering of a bifurcating tip from a time-lapse imaging video of an epithelial Fucci2a expressing ex vivo cultured mammary gland, at two-hour intervals following the appearance of the cleft with the bifurcation angle indicated. **D)** Average bifurcation angles from 10 Fucci2a videos at incremental hours starting from the appearance of the cleft as indicated by color. **E)** Outlines of the epithelial surface rendering of a bifurcating tip at incremental hours from the start of the time-lapse video as indicated by color (ingression of the cleft is indicated by a black arrow and advancement of the daughter tips by white arrows). **F)** The depth of the cleft and the length of the daughter tips were measured from time-lapse imaging videos of surface rendered bifurcating tips after the appearance the cleft and their relative velocity calculated based on duration of the video (n = 10). **G)** Maximum intensity projections of mammary epithelial organoids, derived from R26R-mTmG embryos with (left) or without (right) K14-Cre expression. **H)** Maximum intensity projections of an epithelial organoid cultured and imaged in 3D Matrigel™ for a period of 18 hours. The boxed region follows a bifurcating tip at two-hour intervals. **I)** Quantification of branching events from 26 organoids of 4 time-lapse imaging experiments. Data shown in F represents the median (line) with 25th and 75th percentiles (hinges) plus 1.5× interquartile ranges (whiskers). Statistical significance was assessed with the paired t-test.

Whether tip bifurcation can be accomplished chiefly by epithelial cells or whether the action of mesenchymal cells is necessary, is unclear. Classical tissue recombination experiments suggest that the mesenchyme determines the mode of branching events (Kratochwil 1969). Next, we investigated whether tip bifurcations are possible in the absence of the mesenchymal tissue by examining branching of epithelial organoids. Organoids were cultured for two days for branching to be initiated, and then imaged once per hour by 3D confocal microscopy for a minimum of 19 hours **(Figure 7G)**. Although patterning of the epithelium was clearly atypical, with shorter and more densely packed branches, distinct tip splitting and budding events could be discerned. Quantification revealed that about 40% of the branching events were tip splitting events (**Figure 7H, Video 7**) indicating that tip bifurcation ability is intrinsic to the epithelium.

## Discussion

Branching morphogenesis creates an arborized epithelium by iterating branch elongation and branch point generation in terminal branches of all organs. Although side-branching is prevalent in the adult mammary gland, pubertal branching morphogenesis has been primarily characterized by mesenchymal invasion and bifurcation of the TEBs (Hannezo et al. 2017). How these two actions are executed and integrated at the cellular level, has not been established mainly due to lack of comprehensive real time evidence. Here we have studied branching morphogenesis of the embryonic mammary gland within the native inductive mesenchyme by ex vivo live imaging. We asked how cell cycle activity and cell motility, two basic elements of epithelial construction, are coordinated within bifurcating terminal branches, based on unbiased Fucci2a labeling. We find that directional cell movement gradually increases towards the tip along an elongating branch, which in turn creates a retrograde flow of lagging cells to the trailing duct, sustained by tip cell proliferation. This flow gets interrupted when tips bifurcate, as cell cycle activity and movement is locally repressed at the branch point between nascent daughter tips. Cells within the daughter tips resolve into separate streams of directional cell movement and begin to elongate two new branches.

The branching tips of several mammalian organs are enriched in proliferative cells (Varner & Nelson 2014), but whether differential growth makes branches grow longer or bifurcate, has not been confirmed so far. The proliferative TEBs of the pubertal mammary gland are comprised of cycling stem cells (Scheele et al. 2017). Interestingly, elaborate branching structures can be created solely based on spatially controlled stem cell divisions at the tips of higher plants, in the absence of cell movement (Carles & Fletcher 2003). Our examination of embryonic mammary glands revealed an elevated cell cycle activity within tips, although only in the luminal compartment. Yet cells do not seem to accumulate in the tip, as tip width remained unchanged during branch elongation ex vivo and only slightly increased with branch length in vivo. We found that cells get displaced from the tip based on their differential motility. This may also explain why in pubertal glands some of the tip cell progeny at the border of the TEB are traced to the duct, where they undergo differentiation (Scheele et al. 2017). The progeny of cells at the front part of the TEB were suggested to remain there. Although ductal elongation is apparently fueled by a reservoir of proliferative tip cells, examining the effects of global cell cycle repression let us to conclude that elongation of the duct cannot be based on spatially controlled cell divisions, asymmetric or otherwise. Inhibitor experiments in stroma-free mammary epithelial organoids also point towards this direction (Huebner et al. 2016).

Elongation of embryonic mammary epithelial branches was coupled with directional cell migration, gradually increasing towards the leading edge of the tip. Migration contributes to branching morphogenesis of the vertebrate vasculature and drosophila trachea, where branches elongate upon the invasive migration of “leader cells” occupying the pointed edges of tips and sending out adhesive protrusions (Varner & Nelson 2014). According to our data, a substantial proportion of basal tip cells rearranged into the luminal compartment, suggesting that the leading edge position may be transient. Lack of fixed leader cells was also observed in organoids (Ewald et al. 2008). It is also not evident that leading mammary epithelial cells, encapsulated by the BM, would directly engage the stromal ECM for their motility (Ewald et al. 2012, Dawson et al. 2021) – in fact ECM degradation is needed to allow the TEBs to pass into the stroma in vivo (Feinberg et al. 2018). Stratification seems to be a necessary event for collective cell migration to occur (Ma et al. 2022), which is consistent with the stratification of embryonic branch tips and TEBs. Based on these considerations, branch elongation can be considered to be a process of propulsion, where epithelial cells leverage each other to push the TEB forward (Goodwin & Nelson 2020). We propose that branch elongation is supported by differential cell motility along the branch, coupled with directional cell competition to the leading-edge position. This creates a retrograde flow of lagging cells from tip to duct, consistent with clonal lineage tracing data in pubertal glands (Scheele et al. 2017). Notably, we failed to find evidence of epithelial surface expansion via radial cell intercalation, which contributes to organoid branch elongation (Neumann et al. 2018). Either these cells escape our analysis or cells arrange differently when engaged with their native stroma.

Very little is known of branch point formation in the mammary gland. We now find that it is associated with cell cycle repression and reduced cell motility, in contrast to the behaviors that we observe in the tips of elongating branches. Tip proliferation was not acutely required for new branches to be initiated. Bifurcations however were unstable, and the daughter tips became reabsorbed by the parent tip in the absence of proliferation. Whether on the other hand, cell cycle repression is required at the branch point, remains an open question since we had no means to experimentally force the cells to locally proliferate. It is reasonable that branch points cease to grow once established. In other organs, bifurcations are not associated with local differences in cell cycle activity (Lang et. al. 2021). In the salivary gland however, differential expansion of the inner and outer bud epithelium creates a mechanical instability, which causes the more expansive surface cell sheet to buckle (Wang et al. 2021, Kim et al. 2021). Surface expansion involved subsurface cell division and reinsertion of new surface cells. We did observe a trend of basal-to-luminal cell movement in elongating and bifurcating branches alike, but could not assess whether these cells divided or returned to the surface. Overall, luminal-to-basal cell movements were rare and thus unlikely to contribute to a buckling mechanism. Lineage tracing data indicate that in the pubertal gland, basal and luminal compartments are maintained separately (Van Keymeulen et al. 2011, Davis et al. 2016, Wuidart et al. 2016). A substantial proportion of basal cap cells progenitors does venture into the TEB body, but undergoes apoptosis (Paine et al. 2016, Dawson et al. 2021).

One key aspect in branch point generation is the establishment of a cleft that ultimately changes the geometry of the tip. Initially, the salivary cleft appears as a narrow space between two cells, where cell-cell adhesions become uncoupled (Kadoya & Yamashina et al. 2010). In the embryonic mammary gland, clefts first appeared as shallow indentations that became narrower as they deepened over time, not unlike how lung bifurcations have been described (Schnatwinker & Niswander 2013, Kim et al. 2015). The relative contribution of tip outgrowth/elongation and cleft ingression varied between bifurcations, but was equal on average. Proliferation and outward motility of cells of the daughter tips is likely to be part of the shape change. We explored the possibility that localized contractility would contribute to cleft ingression, as suggested in the lung (Schnatwinker & Niswander 2013), but found no obvious supracellular F-actin structures as evidence of localized concerted contractile activity. However, NMII inhibition abrogated branch point generation in ex vivo cultured mammary glands and also in mesenchyme-free organoid cultures, suggesting that global and/or transient contractility is required for epithelial branching. Actomyosin contractility is known to power multiple cell behaviors, including cell motility that according to the present study, contributes not only to branch elongation but also tip bifurcation in the embryonic mammary gland. If cell motility is abrogated, this may explain why epithelial branches appeared to thicken instead of elongating under NMII inhibition and why branching events were reduced.

We found that cell motility contributes to bifurcations in two ways: while cell immobilization helps to establish branch points, directional cell movement contributes to growth/elongation of the daughter tips. Changes in cell motility were spatiotemporally coordinated with changes in cell cycle activity, implying that they may be controlled by a localized inductive/repressive cue rather than by passive forces. The conditions that might trigger such a cue however are enigmatic, given the variation in branch point frequency in the mammary gland (Lindström et al. 2022). Bifurcations are limited by their distance to the previous branch point, suggesting that a certain minimal tip size or branch length is required (Lindström et al. 2022). Several molecular pathways have been implicated in regulation of branching frequency in the mammary gland (Gjorevski & Nelson 2011, Myllymäki & Mikkola 2019). One possibility is that it involves local modulation of RTK signaling activity, which is a universal regulator of branching morphogenesis (Wang et al. 2017). In the mammary gland, RTK signaling has been associated with tip cell motility and proliferation (Lu et al. 2008, Huebner et al. 2016, Neumann et al. 2018). During kidney bifurcations, cells become sorted in two daughter tips based on Ret activation, which regulates RTK signaling in the kidney (Riccio et al. 2016). Fewer bifurcations are observed when the downstream MAPK pathway is blocked in Mek1/2 double mutant mice (Ihermann-Hella et al. 2014). Branch point generation in the lung has been modeled based on a ligand– receptor-based Turing type of mechanism, involving tissue-restricted expression of the Fgf10–FgfR2b ligand–receptor pair (Lang et al. 2021). How molecular pathways regulate branch point generation is an interesting question for future studies.

In this study, we sought to elucidate the fundamental cellular construction principles of the branching mammary epithelium in its native stroma by imaging ex vivo cultured embryonic mammary glands. Our results suggest that directional cell movement advances the tip of a branch, while differential cell motility and proliferation feeds cells from the tip into the trailing duct during its elongation. Bifurcation of the tip involved local attenuation of both cell proliferation and movement at the branch point while establishment of the daughter tips was facilitated by their sustained proliferation and directional “outward” cell movement. Cleft formation was often, but not always, augmented by “inward” movement of cells in the reverse direction. In a simplified view, our findings imply that branching morphogenesis might be accomplished simply by modulating the same set of cell behaviors in branch tips to alternate between elongation and branch point generation. We observed that tip bifurcations were initiated also in mesenchyme-free cultures, suggesting that they are to some extent, intrinsic to the epithelium. However, stromal collagen deposition is likely to contribute to cleft stabilization and establishment of the bifurcation angle in vivo (Nerger et al. 2021). Future imaging studies are awaited to resolve to what extent the cell behaviors that we have identified here contribute to branching morphogenesis at later stages of development.

## Materials & Methods

### Ethics statement

All mouse experiments were approved by local ethics committee and the National Animal Experiment Board of Finland (licenses KEK19-019, KEK22-014, and ESAVI/2363/04.10.07/2017). Mice were euthanized with CO_2_ followed by cervical dislocation.

### Mouse lines

The conditional R26R-Fucci2a-flox/flox (Mort et al. 2014) reporter mice were maintained in C57Bl/6 background. The STOP cassette surrounded by loxP sites was removed with ubiquitous PGK-Cre (Lallemand et al. 1998) to generate the constitutive R26R-Fucci2a-del/del reporter mice, also maintained in C57Bl/6 background. R26R-Fucci2a-del/del mice were crossed with wild type NMRI mice for cell cycle analysis from fixed samples. R26R-mTmG mice obtained from The Jackson Laboratory (Stock 007576) were in ICR background. R26R-tdTomato-flox/flox mice, also from The Jackson Laboratory (Stock 007914), were maintained in C57Bl/6 background. K14-Cre43 mice (Andl et al. 2004; hereafter K14-Cre) in NMRI background were used for epithelial expression of conditional fluorescent reporters. Mice carrying both K5-rtTA (Diamond et al. 2003) and TetO-Cre (Mucenski et al. 2003) transgenes (K5-rtTA;TetO-Cre) in NMRI background were used for doxycycline-inducible sparse labeling of epithelial cells. Mice were kept in 12 hr light-dark cycles and food and water were available ad libitum.

### Whole-mount staining and imaging

For whole-mount confocal microscopy of fixed samples, E17.5-E18.5 female embryos were used. After removing the head and limbs, a longitudinal incision was made along the midline from the neck towards the groin and two flanks of skin with five pairs of mammary glands were peeled to the sides and cut off. The flanks were first spread on a plastic surface and then submerged and fixed with 4% PFA in PBS o/n at +4⁰C on an orbital shaker, followed by three washes with PBS. For immunofluorescence staining, samples were blocked and permeabilized with 5% goat or donkey serum, 0.3% Triton X-100 in PBS for 1 hours at room temperature. Primary antibodies were prepared in blocking/permeabilization buffer and incubated with samples for 2-3days at +4⁰C. For EpCAM staining, rat anti-mouse CD326 was used as a 1/1000 dilution (552370; BD Biosciences). Rabbit anti-Laminin was used as a 1/1000 dilution (L9393, Sigma Aldrich). After washing the samples four times for 1-2hours in PBS, secondary antibodies and 5µg/ml of Hoechst (H3570; ThermoFisher Scientific) were added in blocking/permeabilization buffer and incubated with the samples for 2-3days at +4⁰C. Secondary antibodies were purchased from Thermo Fisher Scientific and used as a 1/500 dilution. For staining of actin filaments, 10U/ml of Alexa Fluor 568 Phalloidin (A12380; ThermoFisher Scientific) was added to the secondary antibody mixture. Before mounting, samples were washed as described above.

Stained flanks with Hoechst were placed under the Zeiss Lumar microscope equipped with Apolumar S 1.2x objective, epidermis facing down, and mammary glands visualized under the UV light. Individual glands were carefully dissected out together with the surrounding mesenchyme and mounted on slides using Vectashield (Vector Laboratories). To preserve the 3D structure, several layers of sticky tape were used as a spacer between the slide and the coverslip. Mammary glands were imaged either using the Leica SP8 Upright point scanning confocal microscope or Leica TCS SP8 STED inverted microscope using the HC PL APO 20x/0.75 IMM CORR CS2 objective. For cell cycle analysis, 3D image stacks were acquired of the entire mammary gland number 3 using tile imaging with 0.2 µm/px xy resolution and 1-2µm z step size. For closer analysis of terminal branches/tips, images were acquired from mammary glands 1-5 using 0.1µm/px xy resolution and 1µm z step size.

### Ex vivo culture of embryonic mammary glands

Ex vivo culture of embryonic mammary glands was performed according to a previously established protocol (Lan et al. 2022). Left and right flanks of abdominal-thoracic skin containing five pairs of mammary buds were dissected out from E13.5 embryos. The tissues were treated for 20-40 min at +4⁰C with 1,25U/ml of Dispase II (4942078001; Sigma Aldrich) in PBS and 3-4 min at room temperature with a pancreatin-trypsin mixture (2.5mg/ml pancreatin [P3292; Sigma Aldrich] and 22,5mg/ml trypsin in Thyrode’s solution pH 7.4). The enzyme solution was replaced with culture media (10% FBS in 1:1 DMEM/F12 supplemented with 100µg/ml ascorbic acid, 10U/ml penicillin and 10 mg/ml streptomycin) and tissues rested on ice for a minimum of 30min. The epidermal layer was pulled away with small needles, leaving the mesenchymal tissue with the mammary buds. The tissues were collected on square pieces of Nuclepore polycarbonate filter (WHA110605, Whatman) and suspended on metal grids on a 3.5cm Ø plastic dish, filled from below with culture medium. The explants were cultured up to 7 days in +37⁰C with air and 5%CO2, replacing the media every other day. Blebbistatin (B0560, Sigma Aldrich) were added into the culture media as a 20µM concentration on day 5-6 of culture and left for 24 hours. Mitomycin C was added into the culture media as a 1µg/ml concentration, left for two hours, and replaced with normal culture media, where explants were cultured for 22 hours. Branching was assessed by quantifying the number of tips/gland from images taken before treatment and after culture with the Zeiss Lumar microscope equipped with Apolumar S 1.2x objectiveZeiss Lumar xx Fluorescence Stereomicroscope of explants derived from K14-Cre;R26R-mT/mG embryos. EdU labeling was performed by treating the explants with xx µg of EdU in culture media for 1 hours at +37⁰C with 5% CO2 in air atmosphere. Staining of EdU labeled samples was performed with the Click-iT™ EdU Cell Proliferation Kit for Imaging (C10340; Thermo Fisher Scientific) according to the manufacturer’s instructions, with the following modifications.

### Mesenchyme-free mammary rudiment culture

Mammary epithelial rudiments were cultured in 3D Matrigel as previously described (Lan et al. 2022). Skin flanks with mammary rudiments 1-5 were collected from E16.5 female embryos as described above for later stages. Tissues were treated for 1 hour at +4 ⁰C with 2.5U/ml of Dispase II in PBS, followed by 5-7min of treatment with pancreatin-trypsin at room temperature. The tissues were then rested on ice for 30-60min in culture medium. The epidermis was pulled away with needles and mammary rudiments 1-3 liberated from the mesenchyme – at this stage the thoracic mammary rudiments have developed 2-3 small branches. The mesenchymal tissue was cleaned away with needles and the buds were washed by pipetting through a 10µl tip several times. The epithelial rudiments were transferred onto the bottom of a 3.5cm Ø dish with 10µl of culture media. The media was removed and replaced with a 10µl drop of growth-factor reduced Matrigel (356231; Corning), 3-5 mammary rudiments/drop. The rudiments were dispersed as not to touch each other or the bottom of the dish. The Matrigel was allowed to solidify at +37⁰C and when solidified, 2ml of organoid culture media (1:1 DMEM/F12 supplemented with 10U/ml penicillin, 10 000µg/ml streptomycin, 1X ITS Liquid Media Supplement [I3146, Sigma Aldrich] and 2.5nM hFGF2 [CF0291; Sigma Aldrich]). Mammary rudiments were cultured up to 3 days at +37⁰C with 5% CO_2_ in air atmosphere, with the medium replaced on day 2. Blebbistatin was added on day 2 as a 20µM concentration.

### Live imaging

#### Wide-field imaging of branching morphogenesis ex vivo

To monitor the effects of Mitomycin C on branching morphogenesis by live imaging, mammary explants were dissected from E13.5 K14-Cre;R26R-mT/mG embryos and prepared for ex vivo culture as described above, but directly placed with a Pasteur pipette to the apical side of a 24 mm Ø Trans-well insert to be cultured on a 6-well plate (Falcon). Tissue culture medium was added up to the level of the insert (1-1.5ml) and explants cultured for 3 days in a tissue culture incubator at 37⁰C, 5%CO_2_. Imaging was performed using an inverted 3I Marianas wide-field microscope with an environmental chamber (+37⁰C air with 6% CO2) and a long working distance air objective (10x/0.30 N.A. EC Plan-Neofluar Ph1 186 WD = 5.2 M27). A LED light source 183 (CoolLED pE2 with 490 nm/550 nm) was used for exposure and images were acquired at multiple positions using 3-hour intervals over a period of 93 hours between 3-7 days of culture. Between 45-47hrs, samples were treated with 1µg/ml Mitomycin C (MCC) or with the same volume of PBS in fresh normal culture media. After the treatment, media was replaced and imaging continued under normal conditions.

#### Imaging of epithelial rudiments in mesenchyme-free culture

3-5 mammary rudiments, embedded in 10µl drops of Matrigel, were applied onto 3,5cm Ø glass bottom dishes. The organoids were prepared from embryos expressing R26R-mT/mG with or without K14-Cre expression and cultured for 2 days before imaging. The rudiments were imaged at multiple positions with an inverted Leica SP8 Stellaris microscope using the HC PL APO 10x/0.40 CS2 air objective at 1-hour intervals over a period of 19-23 hours. Images stacks were acquired with 1-2µm/px xy resolution using 5-7 µm z steps.

#### Confocal imaging of mammary epithelial cell behaviors ex vivo

Mammary explants from E13.5 K14-Cre;R26R-Fucci2a-flox/flox embryos were established as described above and first cultured 5-6 days in standard culture conditions. Explants were then carefully transferred on Nucleopore filters to the membrane bottom of a 5 cm Ø Lumox dish (94.6077.410; Sarstedt) with the tissue side facing down. The edges of the Nucleopore filter were rapidly attached to the Lumox membrane by touching with fine-tipped forceps dipped in Vetbond tissue adhesive (0200742529; 3M Science), taking care not to touch the explant at the center. A 50µl drop of culture media was added on top of each explant to prevent the tissues from drying. After attaching all the explants, 4-5ml of culture media was added onto the dish. Imaging was performed with Leica SP8 Stellaris or Leica TCS SP8 STED inverted laser scanning microscope with an environmental chamber. The dish was placed on a 3D printed sample holder fashioned for the Lumox dish that can be inserted to the frame of a 6-well plate sample holder on the motorized stage. The environmental chamber was set to +37⁰C air with 5% CO2 and 90% humidity and the samples were equilibrated there for 1 hour before imaging. HC PL APO 20x/0.75 CS2 air objective was used for time-lapse imaging on 4-6 positions at 20min intervals for up to 40 hours. 3D image stacks were acquired at each position with a 0.5-0.8µm/px xy resolution and 2µm z step size, using 600Hz bi-directional scanning. The pinhole was opened to 2AU to reduce the laser exposure needed. For analysis, videos were cropped and time points limited to include only terminal branches that elongated or bifurcated (10-28hrs).

### Fixed image analysis

#### Analysis of whole glands

To analyze E18.5 mammary glands, a surface rendering of the epithelium was first generated using the hand drawing tool on every 3-5 slices. A binary mask was created and exported to Fiji. To measure the width of tips, the line tool was used on the maximum intensity projection of the masked epithelial rendering. In cases of overlap, the mask was cleared in part before the projection was made to reveal the tip to be measured. For cell cycle analysis, the surface rendering was used for masking the Fucci2a signal in the mesenchyme so that epithelial expression could be better visualized. Quantification of Fucci2a expressing cells was performed using the Spots function in Imaris software with 6µm and 7µm spot size for G1/G0 and M/G2/S cells respectively. The spots were made slightly smaller than the measured diameter of the nuclei to ensure recognition neighboring cells. The intensity threshold was manually set for each image to ensure the overlay of spots on the fluorescent nuclei. A binary mask was generated based on the G1/G0 and M/G2/S spots. The mask was used as a template to generate the same number of smaller spots that were spatially more separated. The dimensions of the image were modified to yield cubic voxels. A mask was generated based on the smaller spots and for the epithelial surface rendering. The masked channels were exported to Fiji and the spots were fed to the 3D Objects counter and loaded to the 3D manager of 3D ImageJ Suite (Ollion et al. 2013) as separate regions of interest. The distance of the spots to the surface of the gland was determined by mapping them onto a 3D chamfer distance map, created with Morpholib-J from the masked epithelial rendering (Legland et al. 2016). The data was stratified into outer (≤ 6µm) and central cells (> 6µm). Marker images were created using the point tool to mark the position of tips, ducts and branch points – for ducts the marker was placed squarely between two branch points or between the tip and the last branch point. 3D geodesic distance maps were generated running inside the epithelial rendering from each marker to map the distances of spots. Spots were pooled at a given distance from each marker to calculate the % M/G2/S cells. For tips, cells were pooled at a distance of ≤ 100µm. For ducts and branch points, the distance varied based on the thickness of the branch, between 50 – 100µm. The length of the terminal branches was determined based on measuring the distance of the latest branch point to the tip, using a 3D geodesic distance map generated using markers placed in tips.

#### Analysis of bifurcating tips

Bifurcating tips were analyzed more closely from separate images, where a surface rendering of the epithelium was created either based on EpCAM or laminin staining. The angle of the cleft was triangulated in Imaris using the measurement tool. A binary mask was created based on the surface rendering and another one marking the position of the cleft. The binary channels were exported to Fiji and a maximum intensity projection was created. A straight line was drawn from the front of one daughter tip to the other and the distance of this line to the cleft marker was measured, signifying the depth of the cleft. To analyze cell cycle distribution in the bifurcating tip in Imaris, the surface rendering was cut by the neck and at the root of each daughter tip to create separate volumes for daughter tips and the branch point where the percentage of M/G2/S could be calculated. To quantify the intensity of phalloidin staining, the surface rendering was used for separating epithelial and mesenchymal phalloidin staining into separate channels. Spots were generated to mark the position of the cleft and the daughter tips as well as the neck at both sides. A distance transformation was created emanating from each spot and used as basis for a spherical volume around it of a given distance. The sum of intensities in the epithelium and mesenchyme inside the volume was derived. The number of the corresponding pixels was calculated based on the mask of the epithelial rendering and used for determining mean intensities.

#### Cell shape determination

To analyze cell shape in mammary glands R26R-tdTomato-flox/flox females were mated with K5-rtTA;Tet-O-Cre males. On the 13th day from the plug, females were injected with doxycycline hyclate (D9891-5G; Sigma Aldrich) dissolved in sterile PBS in concentration of 4µg per 1g of mouse body mass. Samples were collected from female embryos at E18.5, fixed, stained with EpCAM and Hoechst as described above, and imaged either using the Leica SP8 Upright confocal microscope with HC PL APO 20x/0.75 IMM CORR CS2 objective with glycerol. For cell shape analysis, 3D image stacks were acquired of the close up mammary gland branches with 0.15µm/px xy resolution and 1.04µm z step size. For image analysis, a surface rendering of the epithelium was generated in Imaris. Surfaces were created based on tdTomato fluorescence using a manually set threshold according to the intensity levels and surface grain size set to 0.3 µm. The position of the cells was determined by relative distance to manually placed landmark. For the tip, cells were pooled at a fixed distance of ≤ 100µm from the landmark, while cells located at a distance > 100 µm were considered as part of the duct. Statistical output of surface rendering was used to analyze cell sphericity.

### Live video analysis

#### Quantification of branching events and branch elongation rates

The time-lapse videos in TIFF format, were opened in Fiji and time series registered with the StackReg plugin (Thevenaz et al. 1998). Branching events were quantified and identified by carefully inspecting the adjacent time points. The length of branches was measured frame to frame using the Segmented Line tool in Fiji.

#### Morphometric analysis of tips

The Fucci2a time-lapse videos were drift corrected in Imaris against the origin of the gland, which is marked by a cluster of intensely fluorescent G1/G0 cells. A surface is created around this cluster and tracking of its center is used for drift correction. The field of view was cropped to facilitate faster analysis of terminal branches. The videos were shortened, when necessary, to focus on analyzing terminal branches exclusively when they elongated. For bifurcating tips, the videos were normalized on the time scale based on the timepoint where the cleft first became apparent. The G1/G0 and M/G2/S channels were merged in Imaris and used as a basis to trace a surface rendering by hand at every 2-3 slices, carefully including all epithelial cells. For morphometric analysis, a mask of the intact epithelial rendering was created and exported to Fiji. The width of the tips was measured with the line tool frame by frame. For bifurcating tips, the length of the tip was also measured from the center of the neck to the cleft and further up to a straight line drawn from one daughter tip to the other – the depth of the cleft being the difference between these two measurements. The angle of the cleft was measured with the Angle tool by tracing along the edges of the cleft towards the daughter tips. The surface rendering was also cut into separate volumes to demarcate different tissue domains. In elongating branches, there was no clear curvature to exactly mark the neck of the tip. Therefore, the tip and the trailing duct were separated by cutting the surface rendering where the M/G2/S cells appeared to be enriched. In bifurcating tips, the curvature of the neck became apparent and could be used as a cutting site. The daughter tips and the branch point could be separated into volumes by cutting diagonally from the cleft towards the neck of the tip.

#### Analysis of Fucci2a spots and tracks

6 and 7µm spots were created in Imaris for G1/G0 and M/G2/S cells, respectively. A distance transformation was created from the intact epithelial surface rendering to determine their distances from the surface of the gland. The % of M/G2/S was calculated inside each volume described above. For cell tracking, the diameter of M/G2/S and G1/G0 nuclei was measured and 8 and 9.3 µm spots were created for G1/G0 and M/G2/S cells, respectively. The maximum distance that a spot may move from one frame to the next was restricted to 8 and 9.3 µm and no gaps were tolerated to ensure that tracking followed the same nuclei. The fidelity of tracking was confirmed by manually tracking a subset of cells. The position of tracks was determined by their mean distance to manually placed tissue landmarks or to the surface of the gland based on distance transformations. The front of the tip in an elongating branch was demarcated by a spot and the distances of tracks referred to it. In bifurcating tips, the position of tracks was determined based on whether they were closer to landmarks placed in daughter tips or the cleft. When indicated, the position of the cells at the start and end of the track were compared either to investigate if they had moved from one volume to another or whether they were closer or further away from a tissue landmark. Parameters of cell movement were compared between differently positioned tracks (Velocity = Track length/Track duration, Net velocity = Track displacement length/Track duration, Straightness = xx). In bifurcating tips, tracks were also compared between different stages of morphogenesis, defined based on morphological measurements. The directionality of the cells was analyzed based on their coordinates at the start and end position of the track as displacement vectors. The directionality was determined as the angle between the displacement vector and a reference vector. For the elongating branch, the displacement vector of the tissue landmark placed at the leading front of the tip served as the reference. For the bifurcating tip, the reference vector was the line drawn from the midpoint of the neck towards the cleft.

## Figure legends

**Figure S1.**
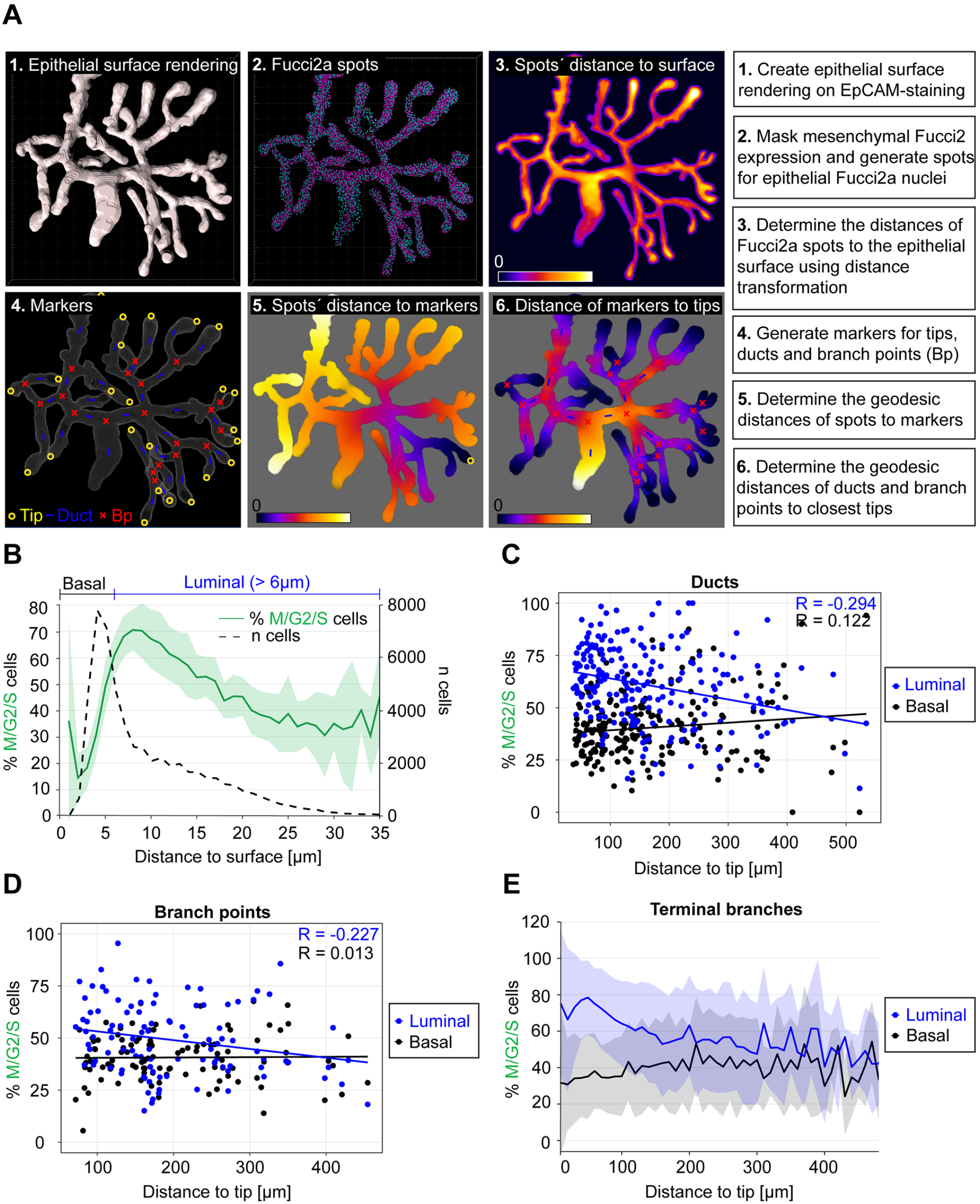
**A)** Pipeline for spatial analysis of cell cycle activity based on the Fucci2a reporter. **B)** Distribution of cell cycle status (% M/G2/S cells) and cell density (n cells) by the distance to the surface of the gland. **C)** Distribution of cell cycle status (% M/G2/S cells) in the “basal” (≤ 6µm) and “luminal” (> 6µm) compartment of ducts (sampled within ≥ 50µm radius of the estimated center point) by the distance to the closest tip. **D)** Distribution of cell cycle status (% M/G2/S cells) in the “basal” and “luminal” compartment of branch points (sampled within ≥ 50µm radius of the estimated center point) by the distance to the closest tip. **E)** Distribution of “basal” and “luminal” cell cycle status (% M/G2/S cells) along terminal branches, from the terminal edge to the branch point.

**Figure S2.**
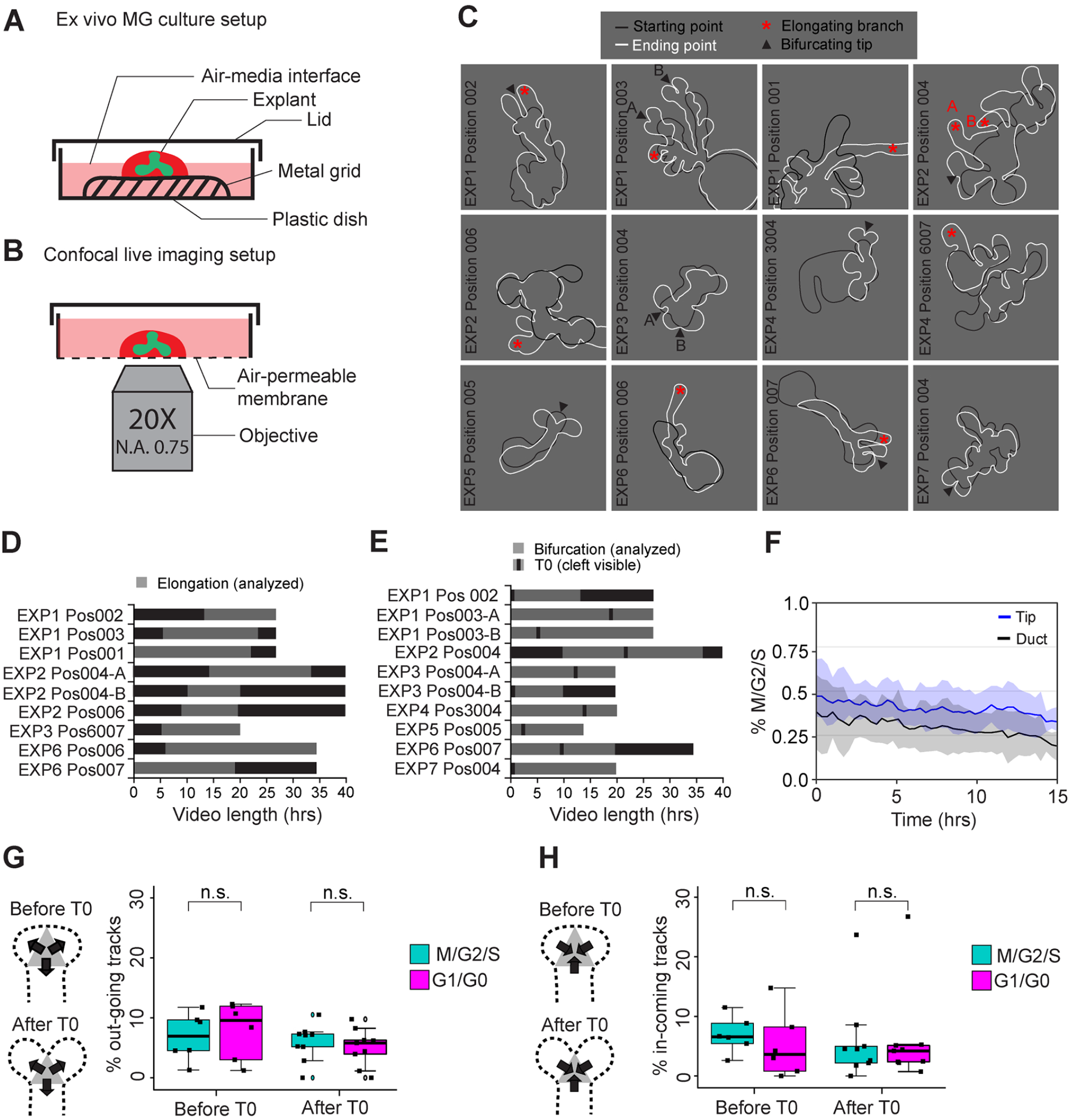
**A)** Depiction of the ex vivo culture setup for the embryonic mammary gland **B)** The picture indicates how explants, first cultured as shown in A, were mounted for confocal live imaging. **C)** Outlines of the ex vivo cultured glands at the beginning (black line) and at the end (white line) of time-lapse imaging, where analyzed elongating terminal branches (marked by a red star) and tip bifurcations (black arrowhead) are indicated. Data was collected from seven imaging sessions and 12 different glands. **D)** The length of videos that were captured, where the period of visible branch elongation was used for analysis (grey). **E)** The length of videos with bifurcating tips (analyzed period in grey), where the timing of different bifurcations was normalized based on the appearance of a cleft (black line; T0). **F)** The effect of long-term time-lapse imaging on the cell cycle activity (% M/G2/S cells) as a function of time, visualized in terminal branches (n = 9). **G)** The percentage of M/G2/S and G1/G0 cells that moved out of the branch point during tip bifurcation (n = 9). **H)** The percentage of M/G2/S and G1/G0 cells that moved into the branch point during tip bifurcation (n= 9). Data shown in G and H represent the median (line) with 25th and 75th percentiles (hinges) plus 1.5× interquartile ranges (whiskers). Data shown in F represent average (line) ± S.D (shaded region). Statistical significance was assessed with the pairwise t-test.

**Figure S3.**
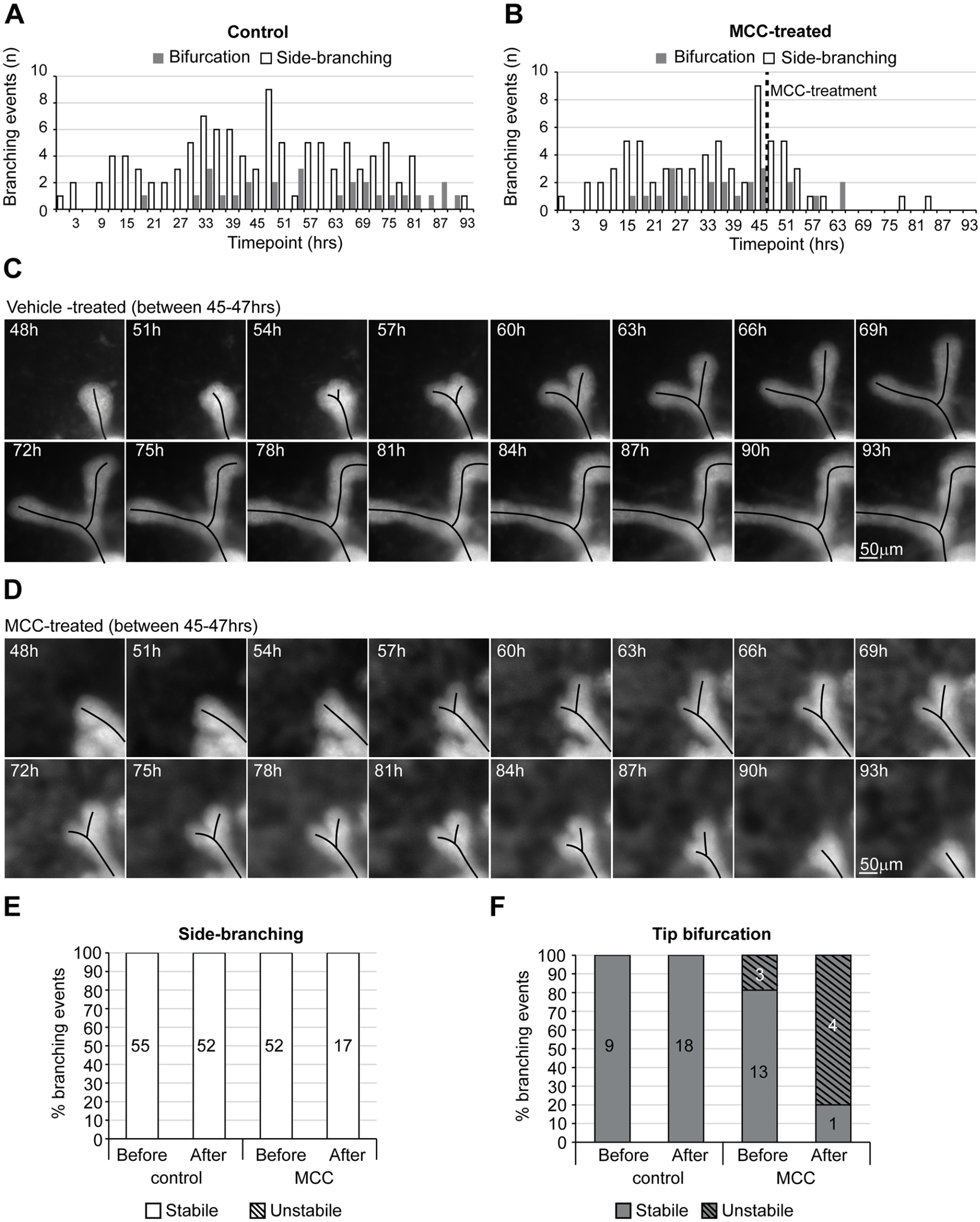
**A)** Occurrence of tip bifurcation and side-branching events in each time point of the time-lapse imaging experiment, where glands were treated with vehicle between 45-47hrs (n^before^ = 64, n^after^ = 70 from 13 glands, two experiments). **B)** Occurrence of tip bifurcation and side-branching events in each time point of the time-lapse imaging experiment, where glands were treated with 1µg/ml of MCC between 45-47hrs (n^before^ = 68, n^after^ = 22 from 12 glands, two experiments). **C)** Captions from a time-lapse imaging video, focusing on a tip bifurcation from a control gland cultured in normal conditions. **D)** Captions from a time-lapse imaging video, focusing on an unstable tip bifurcation that initiated after MCC-treatment. **E)** Side-branching events were evaluated based on whether or not they were maintained once initiated. **F)** Tip bifurcations were evaluated based on whether or not they were maintained once initiated.

**Figure S4.**
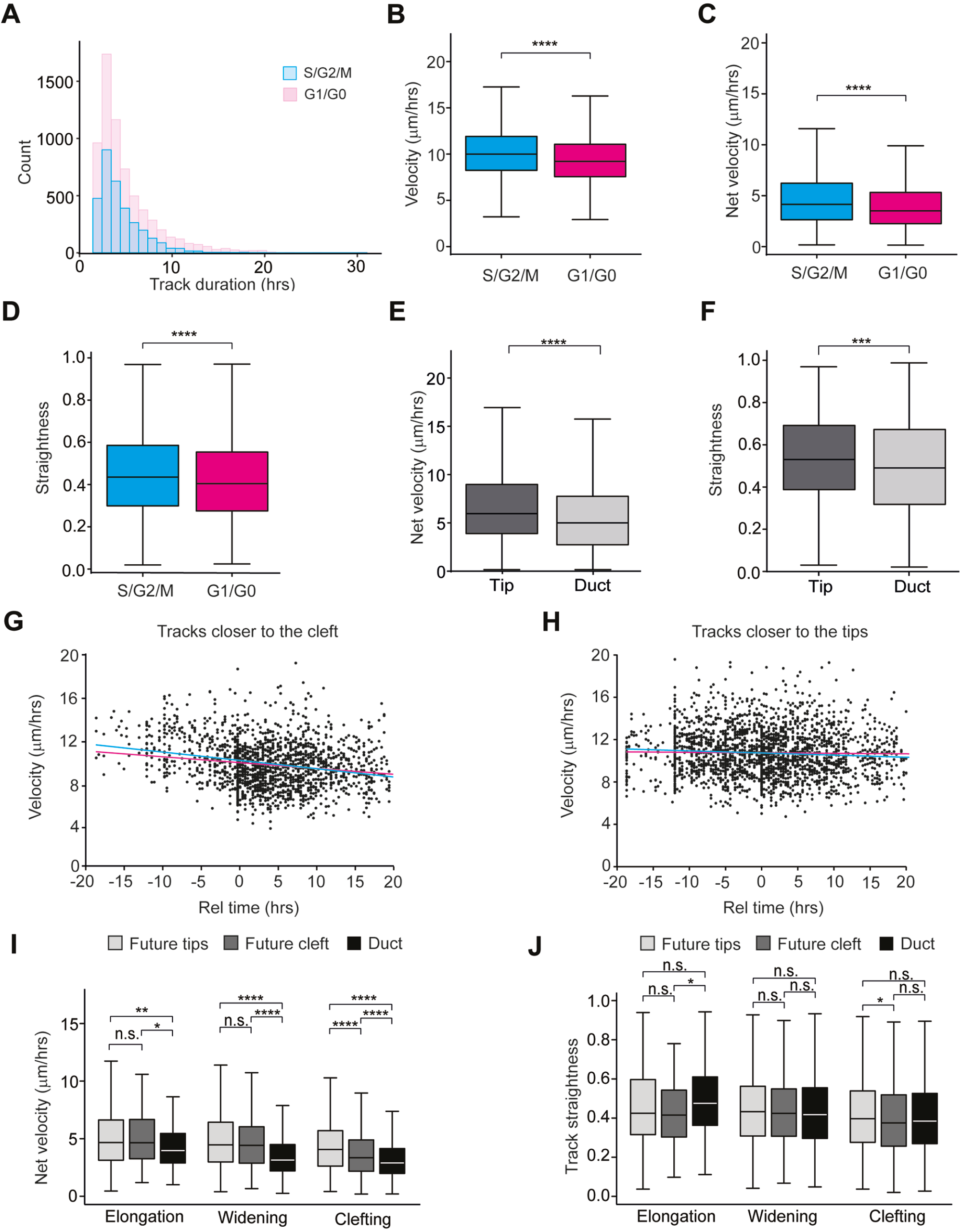
**A)** Histogram of cell tracks of different duration in all time-lapse imaging videos (n^S/G2/M^ = 3191, n^G1/G0^ = 6584). Comparison of the mean velocity **(B)** net velocity **(C)** and straightness **(D)** of tracks of S/G2/M and G1/G0 phase cells (n^S/G2/M^ = 3191, n^G1/G0^ = 6584. Comparison of the net velocity **(E)** and straightness **(F)** of tracks within tip and duct regions of elongating branches (n^tip^ = 600, n^duct^ = 1814). Velocity of tracks within bifurcating tips that were closer to the future cleft **(G)** or closer to future tips **(H)** at different time points relative to the appearance of the cleft at T0 (n^tip^ = 2058, n^cleft^ = 1499 from 10 videos). Net velocity **(I)** and straightness **(J)** of cell tracks within bifurcating tips, divided in those closer to the future cleft (darker grey) and those closer to the future daughter tips (lighter grey), compared those within the duct (black), based on whether they started before bifurcation (n^tip^ = 272, n^cleft^ = 77, n^duct^ = 154), during tip widening (n^tip^ = 688, n^cleft^ = 279, n^duct^ = 1152) or after cleft formation (n^tip^ = 1098, n^cleft^ = 1143, n^duct^ = 1645). Data shown in B-F and I-J represents the median (line) with 25th and 75th percentiles (hinges) plus 1.5× interquartile ranges (whiskers). Statistical significance was assessed with the Student’s t-test, Mann-Whitney test (E & F) and Wilcoxon’s signed rank test (I & J); *, P ≤ 0.05; **, P ≤ 0.01; ***, P ≤ 0.001; ****, P ≤ 0.0001.

## Acknowledgements

We thank Ms. Raija Savolainen and Ms. Riikka Santalahti for excellent technical assistance, and all Mikkola lab members for stimulating discussions. Confocal and widefield microscopy was performed at the Light Microscopy Unit of the HiLIFE-Institute of Biotechnology, University of Helsinki.

This study was financially supported by the Academy of Finland (https://www.aka.fi/en) Center of Excellence Program (grant 307421 to MLM), and project grant (318287 to MLM), Sigrid Jusélius Foundation (http://sigridjuselius.fi/en) (MLM), Jane and Aatos Erkko Foundation (http://jaes.fi/en/) (MLM), Finnish Cancer Foundation (https://www.cancersociety.fi/organisation/cancer-foundation-finland) (MLM), and the HiLIFE Fellow Program (https://www.helsinki.fi/en/helsinki-institute-of-life-science) (MLM); The funders had no role in study design, data collection and analysis, decision to publish, or preparation of the manuscript.

The authors declare no competing financial interests.

